# Duplication of superoxide dismutase and a mutation in aquaglyceroporin mediates the sensitivity of *Plasmodium falciparum* to cryptosporin, a natural product derived from *Acaromyces ingoldii*

**DOI:** 10.64898/2026.06.08.730986

**Authors:** Tiantian Jiang, Jennifer E. Collins, Jin Woo Lee, Sebastian Buss, Basil T. Thommen, Rebecca C.S. Edgar, Karen Wendt, Daisy Wei Chen, Carissa Li, Nimisha Mittal, Raphaella Paes, Natalia Mojica Santos, Leticia Tiburcio Ferreira, Jasveen Bhasin, Jeremiah D. Momper, David A. Fidock, Marcus Lee, Manoj T. Duraisingh, Eric Beitz, Robert H. Cichewicz, Debopam Chakrabarti, Elizabeth A. Winzeler

## Abstract

Cryptosporin, a fungal metabolite, exhibited potent antimalarial activity against both asexual blood stage *Plasmodium falciparum* and liver-stage *Plasmodium berghei* with minimal human HepG2 toxicity. Unlike atovaquone, cryptosporin’s mechanism is independent of mitochondrial electron transport. Minimum inoculum of resistance showed a low risk of resistance development. RNA-Seq analysis revealed the upregulation of genes associated with sexual development including many canonical markers such as Pfs25, and PfCCp3, suggesting a stress response that is also seen when parasites are treated with artemisinin. *In vitro* evolution and whole genome sequencing analysis identified a mutation (F138Y) in PfAQP (PF3D7_1132800) and duplications of the two superoxide dismutase genes, PfSOD-1 (PF3D7_0814900) and PfSOD-2 (PF3D7_0623500). CRISPR/Cas9 editing confirmed that the F138Y mutation in PfAQP was sufficient to confer resistance to cryptosporin. Alignment of the *P. falciparum* structure with that of HsAQP3 suggests the mutation may impact transport of hydrogen peroxide and the transition between open and closed conformations. Indeed, studies with BY4742 Δfps1 yeast expressing PfAQP showed that the permeability of PfAQP was not affected by cryptosporin and that it is likely not a direct target. Taken together, this study highlights the role of PfAQP in the resistance development of cryptosporin. In addition, cryptosporin likely induces high levels of oxidative stress which results in the duplications of oxidative dismutase genes as part of the parasite’s defense response. These findings highlight the role of PfAQP in mediating drug resistance, the mechanism of which warrants further research.

## Introduction

The success of sourcing antimalarials from medicinal plants such as cinchona (quinine) and sweet wormwood (artemisinin) highlights the critical role of natural products in mankind’s battle against diseases. These compounds, including their derivatives (e.g., chloroquine, artemether, artesunate), have profoundly reduced early childhood mortality and saved countless lives (Siqueira-Neto et al., 2023). While natural products have limitations, such as challenges associated with isolation, high production costs, and suboptimal drug-like properties, studying their mechanisms of action can reveal high value targets. Such targets may subsequently be used to discover less toxic, inexpensive, and orally bioavailable drugs. Therefore, the discovery and investigation of new natural product-derived antimalarials remain a critical component in the ongoing fight against malaria.

Cryptosporin was first isolated from *Cryptosporium pinicola* in 1973 and shown to have weak activity against Gram-positive bacteria (Closse & Sigg, 1973). The levorotatory stereoisomer ((-)-cryptosporin) was synthesized in 1989 (Gupta & Frank, 1989). Subsequently, it was independently isolated from various other organisms including *Streptomyces sp.* (Zhang et al., 2013) and *Acaromyces ingoldii* (Gao et al., 2016) from marine sediment. The (+)-cryptosporin used in our study was isolated from *Acaromyces ingoldii* found on loblolly pine bolt from the Kisatchie National Forest, USA (Olatinwo & Fraedrich, 2019). The crude extract of *Acaromyces ingoldii,* which has broad-spectrum antifungal properties, contains three secondary metabolites including cryptosporin, coryoctalactone D and a third compound similar to cryptosporin (Olatinwo & Fraedrich, 2019; Olatinwo et al., 2019).

Here, we report the antimalarial activity of cryptosporin. Cryptosporin demonstrates dual-stage activity in *P. falciparum* blood stage and *P. berghei* liver stage. This compound is an appealing antimalarial tool compound due to its relatively low molecular weight, accessible synthetic routes, and production by a wide range of organisms. However, its mechanism of action remains unknown since its first discovery half a century ago. Understanding its mechanism of action could pave the way for target-based drug discovery using this compound as a validation compound or potentially a lead. In this study, we used *in vitro* evolution and whole genome sequencing to uncover the compound’s mode of action. The data suggests that cryptosporin induces oxidative stress in parasites. Additionally, we find that the resistance mechanism of cryptosporin involves a bi-functional aquaglyceroporin (PF3D7_1132800), which has not been previously associated with drug resistance in *P. falciparum*.

## RESULTS

### Natural product cryptosporin demonstrates antiplasmodial activity in blood and liver stages

In an effort to identify natural products with antiplasmodial potential, a library of 677 pure compounds isolated from fungi and other natural product sources had previously been screened at a fixed concentration of 1 µM with subsequent inhibition in dose response format determined in the multidrug resistant *P. falciparum* strain, Dd2 (Collins, 2022). These efforts had led to the discovery of 57 compounds with an EC_50_ < 1 µM, including the fungal secondary metabolite cryptosporin (Figure 1A).

**Figure 1.**
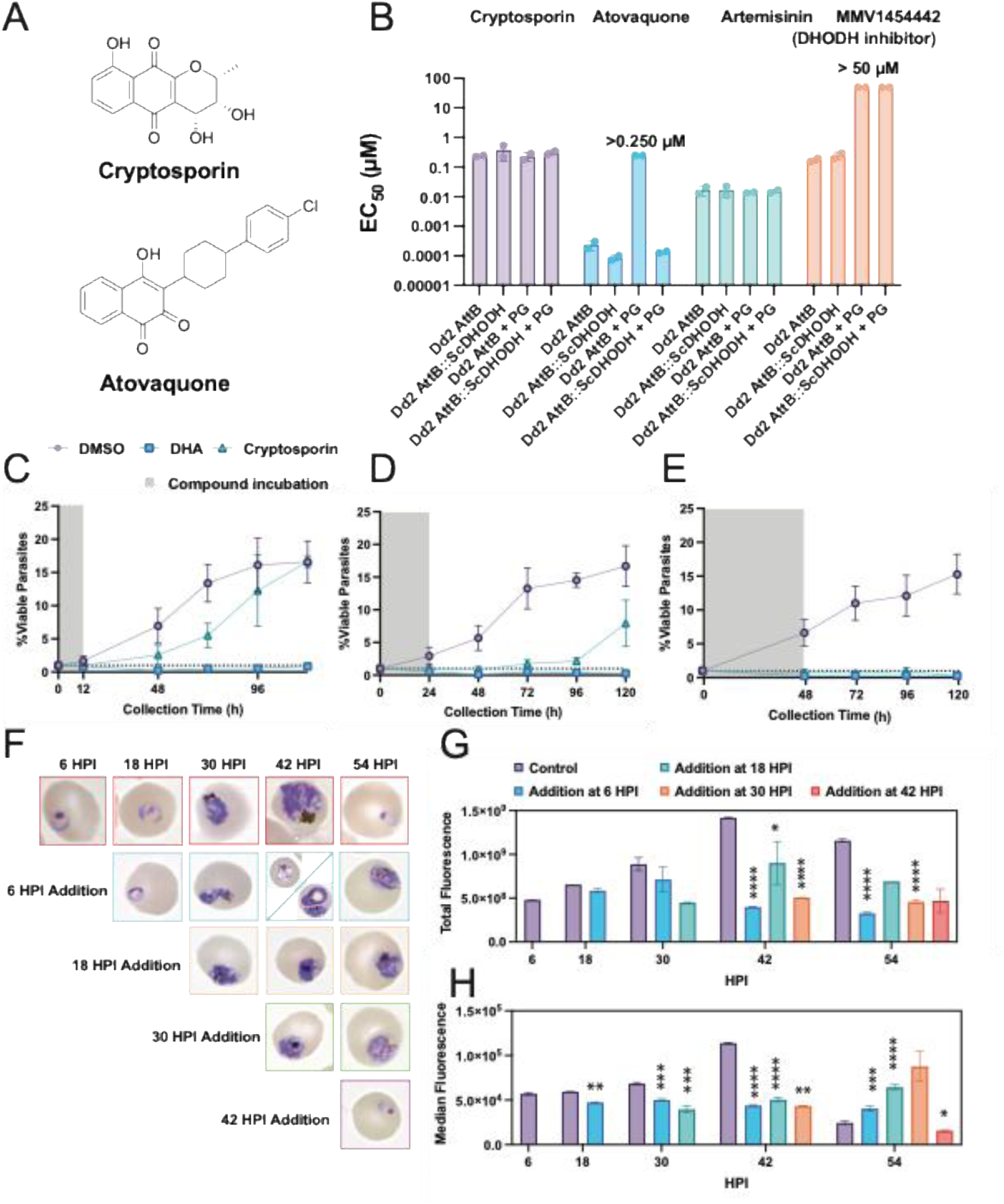
Antiplasmodial killing profile of cryptosporin and its stage specific inhibition. **(A)**The structures of cryptosporin and atovaquone. Both compounds contain a naphthoquinone scaffold **(B)** The mode of action of cryptosporin is not related to DHODH or mitochondrial ETC. Cryptosporin and the control compound atovaquone were screened against two lines (Dd2 attB and Dd2 attB::ScDHODH) with or without proguanil (PG). Dd2 attB indicates the parental line, whereas Dd2 attB::ScDHODH is the parasite line expressing yeast DHODH. Data are from two biological replicates with four technical replicates. **(C, D, E)** Growth curves of *P. falciparum* Dd2 parasites, following exposure to a 10 × EC_50_ concentration of cryptosporin (green), fast acting dihydroartemisinin (DHA) (blue), or DMSO vehicle control (purple) for varying incubation times (12h(**C**), 24h(**D**), or 48h(**E**)) as indicated in grey. Viable parasitemia was determined by flow cytometry using MitoTracker Red and SYBR Green I. Gating was done using DHA and vehicle controls, as well as staining and RBC-only controls. **(F,G,H)** Stage specific growth inhibition of cryptosporin. Synchronous Dd2 parasites were treated with vehicle control DMSO or a 5 × EC_50_ concentration of cryptosporin at select time points 6h, 18h, 30h, or 42 h post-invasion (HPI). Culture progression was then monitored every 12 h via flow cytometric histogram analysis with YOYO-1, and thin smear Giemsa stain. **(F)** Giemsa staining of parasites treated with cryptosporin or DMSO vehicle control at selected collection timepoints. **(G)** Total fluorescence signal across replicates. **(H)** Median fluorescence signal across replicates. Statistical significance for the 18, 30, and 42 HPI treatments was determined by two-way ANOVA followed by Šídák’s multiple comparisons test to compare treatment groups across timepoints (****p<0.0001; ***p<0.001; **p<0.01, *p<0.05). As there was only one timepoint for the 42 HPI treatment (collection at 54 HPI), a paired t-test relative to the vehicle control was used to determine significance for this point. Results are representative of two biological replicates.

We further characterized the antiparasitic properties of cryptosporin which showed an EC_50_ of 0.57 µM in Dd2 line (**Table 1**). Antiplasmodial activity was also evaluated in the drug-sensitive 3D7 strain, where a comparable EC_50_ value of 0.89 μM was observed, corresponding to a resistance index of 1.6. Given the increased need for inhibitors with multistage antiplasmodial activity, cryptosporin was then assayed against the plasmodial exoerythrocytic (liver) stage. To this end, sporozoites from a luciferase-expressing line of *P. berghei* (*Pb*Luc) were added to human HepG2 cells, with compound added pre-invasion. Both parasite inhibition and human cell toxicity were then determined. Strikingly, cryptosporin showed near equal levels of inhibition against *P. berghei* liver stages compared with *P. falciparum* asexual blood stage with a *Pb*Luc EC_50_ of 0.64 µM. Cytotoxicity assay in HepG2 cells showed an EC_50_ of 6.03 µM, which equates to a selectivity index of 9.4. To determine whether cryptosporin is susceptible to known antimalarial resistance mechanisms, we tested its activity against mutant lines PfPI4K_S1320L (McNamara et al., 2013), PfCARL_I1139K (LaMonte et al., 2016), and PfACS_A597V (Summers et al., 2022). We did not observe significant cross-resistance, suggesting a distinct mechanism of action (**Table 1**).

**Table 1.** Antiplasmodial activity of cryptosporin.

| Asexual Blood Stage<sup>a</sup> |  | Extraerythrocytic Stage<sup>b</sup> |  | Asexual Blood Stage<sup>b</sup> |  |  |  |  |  |
| --- | --- | --- | --- | --- | --- | --- | --- | --- | --- |
| Parasite Line | EC<sub>50</sub> (μM) | Cell/Parasite Line | EC<sub>50</sub> (μM) | Parasite Line | EC<sub>50</sub> (μM) | Parasite Line | EC<sub>50</sub> (μM) | Parasite Line | EC<sub>50</sub> (μM) |
| *Pf*Dd2 | 0.57 ± 0.02 | *Pb*Luc | 0.64 ± 0.30 | *Pf*Dd2 | 0.83 ± 0.1 | - | - | - | - |
| *Pf*3D7 | 0.89 ± 0.11 | HepG2 | 6.0 ± 0.04 | *Pf*PI4K (S1320L) | 0.70 ± 0.05 | *Pf*CARL (I1139K) | 1.0 ± 0.01 | *Pf*ACS (A597V) | 0.97 ± 0.05 |
| RI<sup>c</sup>: | 1.6 | SI<sup>d</sup>: | 9.4 | RI: | 0.84 | RI: | 1.2 | RI: | 1.2 |

<sup>a</sup>Results represent the mean and SEM of 3 biological replicates using Chakrabarti lab lines.
<sup>b</sup>For the asexual blood stage assays, results represent the mean and SEM of 2 biological replicates with 3 or 4 technical replicates using Winzeler lab mutant lines (*Pf*PI4K\_S1320L (McNamara et al., 2013); *Pf*CARL\_I1139K (LaMonte et al., 2016); *Pf*ACS\_A597V (Summers et al., 2022)). Extraerythrocytic stage EC<sub>50</sub>s were conducted with 4 biological replicates and 4 technical replicates.
<sup>c</sup>RI: resistance index. Calculated as EC<sub>50</sub> of mutant line over EC<sub>50</sub> of wild-type line.
<sup>d</sup>SI: selectivity Index. Calculated as EC<sub>50</sub> in the HepG2 over EC<sub>50</sub> in *Pb*Luc.

To determine if cryptosporin would be suitable for *in vivo* antimalarial testing, we evaluated the physicochemical and pharmacokinetic properties of cryptosporin, which are summarized in **Table 2**. Cryptosporin exhibited a kinetic solubility of 176 µM and 57.7% of the compound remained after 60 minutes in mouse liver microsomes. Following intraperitoneal administration at 20 mg/kg in mice (n = 4), cryptosporin reached a peak plasma concentration (C_max_) of 1640 ± 276 ng/ml at 30min. The area under the concentration–time curve (AUC_last) was 4229 ± 1153 hr·ng·mL⁻¹. The apparent volume of distribution (Vz/F) and clearance (CL/F) were 4.79 ± 1.06 L·h⁻¹·kg⁻¹ and 32.0 ± 8.61 L/kg, respectively. Taken together, these data showed that cryptosporin has favorable solubility and moderate stability with fast absorption. However, its *in vivo* exposure profile may limit its utility without further optimization.

**Table 2.**
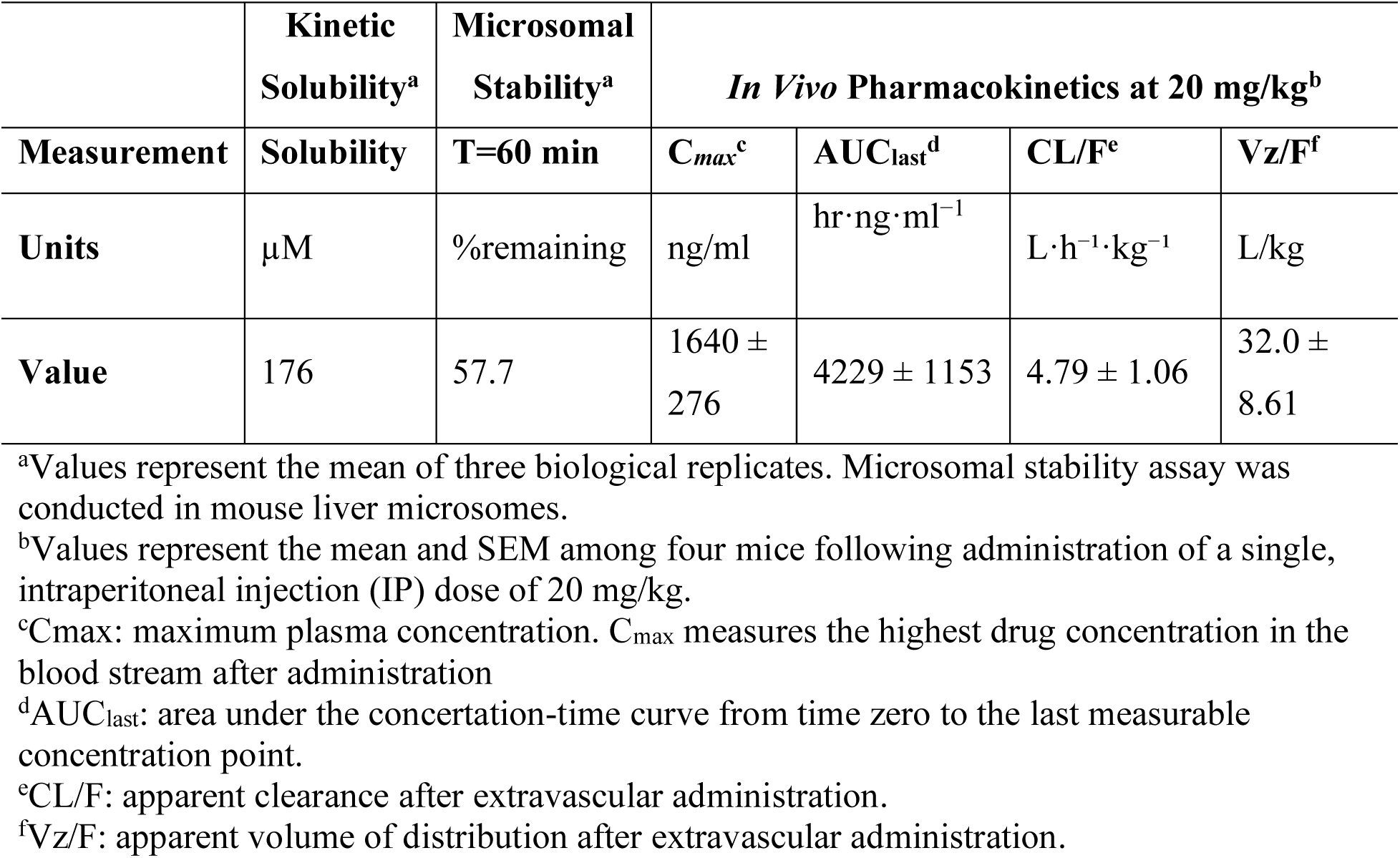
Physiochemical and pharmacokinetic properties of cryptosporin.

|  | Kinetic Solubility<sup>a</sup> | Microsomal Stability<sup>a</sup> | In Vivo Pharmacokinetics at 20 mg/kg<sup>b</sup> |  |  |  |
| --- | --- | --- | --- | --- | --- | --- |
| Measurement | Solubility | T=60 min | C<sub>max</sub><sup>c</sup> | AUC<sub>last</sub><sup>d</sup> | CL/F<sup>e</sup> | V<sub>z</sub>/F<sup>f</sup> |
| Units | μM | %remaining | ng/ml | hr·ng·ml<sup>-1</sup> | L·h<sup>-1</sup>·kg<sup>-1</sup> | L/kg |
| Value | 176 | 57.7 | 1640 ± 276 | 4229 ± 1153 | 4.79 ± 1.06 | 32.0 ± 8.61 |

<sup>a</sup>Values represent the mean of three biological replicates. Microsomal stability assay was conducted in mouse liver microsomes.
<sup>b</sup>Values represent the mean and SEM among four mice following administration of a single, intraperitoneal injection (IP) dose of 20 mg/kg.
<sup>c</sup>C<sub>max</sub>: maximum plasma concentration. C<sub>max</sub> measures the highest drug concentration in the blood stream after administration
<sup>d</sup>AUC<sub>last</sub>: area under the concentration-time curve from time zero to the last measurable concentration point.
<sup>e</sup>CL/F: apparent clearance after extravascular administration.
<sup>f</sup>V<sub>z</sub>/F: apparent volume of distribution after extravascular administration.

### Cryptosporin acts independent of the mitochondrial transport chain

We next to investigate its mode of action and other parasitological characteristics. Like the known antimalarial atovaquone, cryptosporin contains a naphthoquinone scaffold (**Figure 1A**). As such, we sought to determine whether cryptosporin functions similarly to atovaquone through the inhibition of mitochondrial electron transport chain (ETC). To this end, the EC_50_ of cryptosporin was verified in a transgenic line that expresses *Saccharomyces cerevisiae* dihydroorotate dehydrogenase (ScDHODH). Expression of ScDHODH, a structurally distinct cytoplasmic enzyme that uses fumarate instead of quinone as the electron acceptor, can rescue *P. falciparum* parasites treated with ETC inhibitors (Painter et al., 2007). Inhibitors of DHODH (e.g. DSM265) and mitochondrial function (e.g. atovaquone) can be further distinguished by co-treatment with proguanil (PG), which acts synergistically only with ETC inhibitors, thus restoring drug sensitivity (Painter et al., 2007). Here we used a parasite line in which ScDHODH was integrated into an introduced attB locus via the Bxb1 integrase system (Ke et al., 2011; Nkrumah et al., 2006). Testing with the transgenic line (Dd2 attB::ScDHODH), as well as its isogenic parent (Dd2 attB) showed cryptosporin was equally potent in both lines with or without PG (Table 1, Figure 1A), while the control drug atovaquone (a cytochrome bc1 inhibitor) showed a synergistic effect with PG in the isogenic parent line (Dd2 attB) (**Figure 1B**). Atovaquone was not potent (>250nM) in the ScDHODH-expressing transgenic line but treatment with PG restored its potency (**Figure 1B**). These data indicate that cryptosporin does not act via DHODH nor the mitochondrial ETC and suggests a novel mechanism of action.

### Cryptosporin demonstrates a low resistance risk and a moderate killing profile

We next sought to further characterize the compound by assessing its rate of kill as well as the potential for parasites to acquire resistance to cryptosporin. To determine the resistance risk, a minimum inoculum of resistance (MIR) assay was conducted in which drug pressure at 3 × IC_90_ was applied to a clonal Dd2-B2 line and then recrudescence was monitored for 60 days. Selections were applied to 12 wells at 1.2 × 10^6^ parasites per well and three wells at 1.0 × 10^7^ parasites per well, for a total of 4.4 × 10^7^ parasites. No parasite recrudescence was seen up to 60 days. This suggests that the MIR of cryptosporin is greater than the starting inoculum of 4.4 × 10^7^, placing it at a lower risk of developing resistance (Duffey et al., 2021).

To further define the antiplasmodial activity of cryptosporin, its *in vitro* killing activity was profiled following varying incubation times. Cryptosporin was added at a 10 × EC_50_ concentration to asynchronous Dd2 parasites and incubated for 12h, 24h, or 48 h, alongside a vehicle or fast-acting DHA control. After compound removal, cultures were monitored every 24 h using flow cytometry staining with SYBR Green I and MitoTracker Red (MTR) to assess the percentage of viable parasites (SYBR positive, MTR positive). In the 12 h incubation condition, cryptosporin showed an immediate effect on parasite reduction compared to the vehicle control (**Figure 1C**). However, the parasitemia recovered, reaching a similar level to the control by day 5. With 24 h incubation, cryptosporin demonstrated a marked improvement to inhibition, with parasitemia not increasing until day 3, and a substantial decrease in viable parasites on day 5 compared to the control (**Figure 1D**). The extended 48 h incubation with cryptosporin resulted in complete inhibition for all 5 days, with no notable change to the percentage of viable parasites (**Figure 1E**). Taken together, these results suggest cryptosporin shows a moderate killing profile, having an immediate effect on parasitemia but requiring more than 24 h of incubation for full inhibition.

### Incubation with cryptosporin induces growth arrest at the late trophozoite stage

To define the stage at which cryptosporin exerts its antimalarial activity and to gain insight into its mechanism of action, we performed a stage-specific inhibition assay across the intraerythrocytic developmental cycle. Synchronized Dd2 parasites were treated with 5 × EC_50_ cryptosporin or vehicle control DMSO starting at 6h, 18h, 30h, or 42 h post-invasion (HPI). Parasite growth was monitored by flow cytometry and microscopy. Samples were collected every 12 hours until reinvasion (54 HPI) for Giemsa staining and, in parallel, stained with YOYO-1 and analyzed by flow cytometry. Incubation with cryptosporin at all timepoints prior to late schizogony (42 HPI) resulted in life cycle inhibition at the late trophozoite stage, although minor variations were observed (**Figure 1F-H**). In all cases (6 to 30 HPI addition), there was a reduction in DNA accumulation (as measured by median fluorescence intensity) and no segmentation was observed (**Figure 1F, 1H**). No reinvasion was seen at 54 HPI and parasites appeared smaller with irregular hemozoin structures and localization (**Figure 1F**). Conversely, parasites collected following the final compound addition at 42 HPI were able to reinvade and appeared structurally normal with a noticeable but not statistically significant decrease in total fluorescence (**Figure 1F-H**).

### Transcriptome analysis shows induction of gametocytogenesis after cryptosporin treatment

We next utilized RNASeq to investigate cryptosporin’s impact on parasites. When attempting to harvest RNA from synchronous 3D7 culture treated for 2 or 4 h at 1× EC_50_ concentration, we noted that incubation with cryptosporin led to a greatly reduced RNA yield compared to DMSO vehicle control. As a result, a shorter, one hour incubation was required to obtain the necessary quantity of RNA from treated cultures. Even with this short interval, significantly less total RNA was harvested from an equal volume of culture following cryptosporin treatment, suggesting some impact to RNA storage, production, or extraction efficiency (**Figure S1A**). While on average, no significant difference was seen with the RNA integrity number (RIN) and rRNA ratio (28s /18s) between cryptosporin and vehicle treated culture (Figure S1B-C), one replicate with a low RIN was removed as a precaution. Ribosomal RNAs were depleted and then remaining RNA was reverse transcribed and labeled with random priming. The average read coverage was 261×. After rRNA depletion and library-size normalization, 5,573 genes (protein-coding and noncoding) were considered expressed above background (**Table S1**). We observed good correlation between samples (Spearman rho = 0.94-0.98) (**Figure S2A**). The data showed high level of expression for the usual candidates with genes encoding histones (PF3D7_0610400 (H3), PF3D7_0617800 (H2A) and PF3D7_1105100 (H2B)) showing the most abundant transcripts for protein coding genes.

We next sought to identify differentially expressed genes. Data were first normalized, and 211 genes were identified that showed a two-fold average increase in the cryptosporin set relative to the control set. Interestingly, inspecting this list showed that many cryptosporin-upregulated genes have an annotated role in sexual or transmission stages of the parasite lifecycle. These included a number of well annotated sexual development genes (Table S1) such as Pfs25 (ookinete surface protein P25, PF3D7_1031000, fold change = 2.27, p<0.05) (Schneider et al., 2015), dynactin subunit 6, putative (PF3D7_0818300, fold change = 2.13, p<0.05) (Kengne-Ouafo et al., 2023), male development protein MD5, putative (PF3D7_0414500, fold change = 2.05, p<0.05) (Russell et al., 2023), CCp3 (LCCL domain-containing protein,PF3D7_1407000, fold change= 2.19, p<0.05) (Simon et al., 2009) and NTH (PF3D7_1453500, fold change = 2.39, p<0.05). The latter encodes a membrane-bound NAD(P) transhydrogenase (NTH) found in the apicoplast and crystalloid, a critical enzyme for sporozoite development during the mosquito stage (Saeed et al., 2020). Independent RT-PCR confirmed the upregulation of NTH (**Figure S2B**). We observed few significant metabolic pathway enrichments after correcting for multiple hypothesis testing. Similar patterns have recently been observed after Single-Cell transcriptome analysis after artemisinin treatment, indicating that this may be a general stress response (Godinez-Macias et al., 2026).

To determine whether upregulation during gametocytogenesis is also observed for less-characterized upregulated genes, we examined lifecycle expression using an archived transcriptional dataset with samples from different lifecycle conditions with a focus on gametocytogenesis (Young et al., 2005). Plotting and clustering the lifecycle expression of 165 genes (**Table S2**), which showed ≥ 2-fold upregulation after cryptosporin treatment and were expressed above background in the earlier dataset, revealed that two-thirds of these genes (cluster 1; 102 genes; **Table S2**) exhibited minimal expression during the asexual blood stage but were strongly upregulated during sexual development (**Figure S3**). Gene Ontology (GO) enrichment analysis showed few matches for the 102 genes except for cellular components associated with sexual development, such as the crystalloid (GO:0044312, P = 9.57e-6). A small subset of eight genes was also upregulated in sporozoites (cluster 3) and included sporozoite invasion-associated protein 2 (PF3D7_0830300). Transcripts detected in blood stages (cluster 4, 42 genes, **Table S2**) were strongly enriched for subtelomeric multigene family members involved in antigenic variation (GO:0020033, P = 4.69 × 10⁻⁹), primarily rifins, most of which were expressed near background in the RNASeq data. Overall, these data suggest that cryptosporin treatment either induces gametocytogenesis, which is well known to be induced by stress (Subramaniam et al., 2025) or that sexually committed parasites are protected from cryptosporin treatment.

### *In vitro* evolution experiment with *Plasmodium falciparum* Dd2 identifies mutations that confer resistance to cryptosporin

To further investigate the compound’s mechanism of action*, in vitro* evolution of parasites under stepwise increasing concentration of cryptosporin (from 40nM to 1.2 µM) was performed over 10 months. Parasites were continuously cultured in three parallel T75 flasks as three independent biological replicates. An average of 2-fold EC_50_ shift was first observed five months into the resistance selection and maintained at the end of the selection. At the end of the selection (10 months post-selection), two clones from each flask were selected and subjected to EC_50_ dose response assays. As shown in **Table 3** and **Figure 2A**, an average 2-fold EC_50_ shift was observed in all clones compared to the parent Dd2. Given the evidence of oxidative stress induced by this compound, the six clones were evaluated for cross-sensitivity to artemisinin, a drug whose mechanism of action involves oxidative stress (**Figure 2B**). Only one clone from the third flask (3F5) showed a significant (**p<0.01), 1.8-fold (56.4nM/30.6nM) EC_50_ shift. Selected clonal lines were tested for cross-resistance to atovaquone, no significant EC_50_ shift was observed (**Figure 2C**).

**Figure 2.**
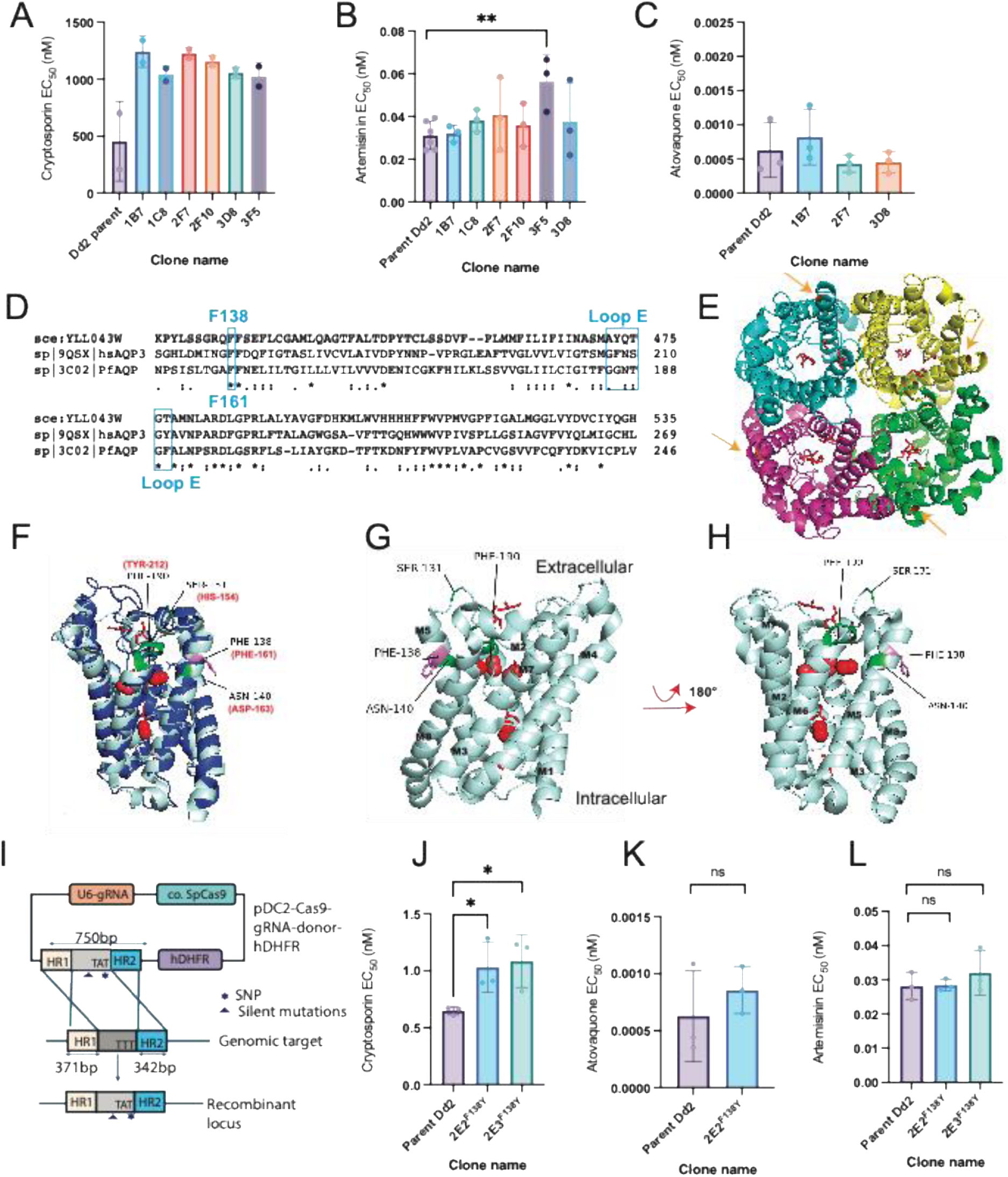
The sensitivity of *in vitro* evolved parasite clones and clones with F138Y introduced via CRISPR/Cas9 to cryptosporin, atovaquone and artemisinin. (**A**) *In vitro* evolved parasite clones showed an EC_50_ difference compared to the parent Dd2 line. (**B**) Among the in vitro evolved parasite clones, 3F5 showed a significant (**p<0.01), 1.8-fold (56.4nM/30.6nM). (**C**) Dose response assay was conducted with atovaquone in selected clonal lines. No significant EC_50_ difference was observed. (**D**) In vitro evolution and whole genome sequencing identified F138Y in PfAQP. Sequence alignment (Clustal Omega) of orthologous aquaglyceroporins from *Saccharomyces cerevisiae* (Scer: YLL043W), *Plasmodium falciparum* (PfAQP: 3C02) and *Homo sapiens* (hsAQP3: 9QSX) shows that F138 is conserved across species. GGNTGF is labeled which corresponds to hsAQP3 GFNSGY (loop E). (**E**) F138Y mutation is highlighted in orange in the tetramer form of PfAQP (PDB: 3C02) seen from the extracellular side. Five glycerol molecules (red sticks) occupy the transduction channel in each monomer. (**F**) Alignment of hsAQP3(PDB: 9QSX) with PfAQP (PDB: 3C02) using PyMOL (Version 2.5.8, Schrödinger, LLC) shows that the phenylalanine at reside 138(magenta) is conserved. The four hydrogen peroxide molecules (red balls) are shown in addition to the five glycerol molecules (red sticks). The residues (hsAQP3 in red, the corresponding PfAQP residues in black) contributing to the close conformation of the pore of hsAQP3 are labeled (Huang et al., 2025). The loop E is highlighted in green. (**G, H**) F138 is located near the hydrogen peroxide channel. (**H**) is a result of clockwise rotation of (**G**) by 180 degrees. Five glycerol molecules delineate the transduction channel. The four hydrogen peroxide molecules were superimposed onto PfAQP from aligning with hsAPQ3. (**I**) Schematic of CRISPR/Cas9 to introduce F138Y in wild type Dd2 parent line. Two clones (2E2^F138Y^ and 2E3^F138Y^) were generated through CRISPR/Cas9 and limiting dilution. (**J, K, L**) The sensitivity of CRISPR/Cas9 edited clones to cryptosporin (**J**), atovaquone (**K**) and artemisinin (**L**). Both clones showed a significant (*p<0.05) EC_50_ shift from parent Dd2 (**J**). No significant EC_50_ shift was observed in atovaquone (**K**) or artemisinin (**L**).

**Table 3.** The phenotype and genotype of clones derived from *in vitro* evolution and CRISPR/Cas9 genetic editing.

| Strain or Clone | EC<sub>50</sub>(μM)<br>(mean± SEM) | Fold change | PfAQP F138Y<sup>c</sup> | PfSOD-1<sup>d</sup><br>Copy Number | PfSOD-2<sup>e</sup><br>Copy Number | Atovaquone EC<sub>50</sub>(nM)<br>(mean± SEM)<sup>f</sup> |
| --- | --- | --- | --- | --- | --- | --- |
| Dd2<sup>a</sup> | 0.58 ± 0.1 | NA | NA | 1 | 1 | 0.63 ± 0.2 |
| 1-B7<sup>a</sup> | 1.2 ± 0.1 | 2.1 | Y | >2 | >2 | 0.82 ± 0.2 |
| 1-C8<sup>a</sup> | 1.1 ± 0.002 | 1.9 | Y | 1 | >2 | NA |
| 2-F7<sup>a</sup> | 1.1 ± 0.1 | 1.9 | Y | >2 | >2 | 0.43 ± 0.1 |
| 2-F10<sup>a</sup> | 1.2 ± 0.04 | 2.1 | Y (61%) | >2 | >2 | NA |
| 3-D8<sup>a</sup> | 1.1 ± 0.05 | 1.9 | Y | 1 | >2 | 0.45 ± 0.1 |
| 3-F5<sup>a</sup> | 1.0 ± 0.09 | 1.7 | Y | 1 | >2 | NA |
| <sup>b</sup>2E2<sup>F138Y</sup> | 1.0 ± 0.1 | 1.7 | Y | 1 | 1 | 0.86 ± 0.1 |
| <sup>b</sup>2E3<sup>F138Y</sup> | 1.1 ± 0.1 | 1.9 | Y | 1 | 1 | NA |

<sup>a</sup>Clones isolated through limiting dilution from *in vitro* evolved parasite lines. The number to the left of the hyphen indicates the flask (No.1,2,3) from which the clone was isolated.
<sup>b</sup>Two clones isolated through limiting dilution from CRISPR/Cas9 edited bulk culture.
<sup>c</sup>The gene ID for PfAQP is PF3D7\_1132800. The presence of the mutation is indicated as “Y” and otherwise it is indicated as “NA.” The allele frequency is shown in parentheses; if not specified, it is 100%.
<sup>d</sup>The gene ID for PfSOD-1 is PF3D7\_0814900.
<sup>e</sup>The gene ID for PfSOD-2 is PF3D7\_0623500.
<sup>f</sup>Dose response assays were performed for selected lines and “NA” indicates that the assay was not performed.

To determine the genetic basis of resistance in these clones, whole-genome sequencing analysis was performed on all six clones (**Table 3**, **S3**; BioProject PRJNA1451291). The reads were aligned to the 3D7 reference genome and variant calling was performed. Newly emerged SNVs were identified subtracting those not found in the isogenic parent sequences (**Table S3**). Sixteen new SNVs were detected relative to the isogenic parent, of which only one was missense and located in the core genome with the mutation appeared in all clones with high frequency (100% except clone 2F10). This missense mutation was predicted to encode an F138Y transition in the aquaglyceroporin gene PfAQP, (PF3D7_1132800). This mutation has been consistently found in previous bulk culture sequencing events (5, 7, and 8 months-post selection, **Table S4**; BioProject PRJNA1451291). The F138 residue (corresponding to F161 in human AQP3) is conserved among orthologous AQPs from human and yeast (**Figure 2D**, **Table 3**). The mutation (F138Y) in aquaglyceroporin is specific to cryptosporin and has not been previously observed in thousands of similarly evolved lines we or the community has created for other compounds (Luth et. al.). Indeed, examination of a databased that consists of our evolved clones as well as published data shows that no evolved missense mutations have been detected in PfAQP after exposure to more than 150 different compounds.

The F138Y mutation was mapped to the 3D structure of PfAQP (PDB: 3C02) with glycerol bound (Newby et al., 2008). PfAQP is a member of the OG7_0001174 (OrthoMCL 7) ortholog group that includes the four human aquaglyceroporin isoforms hsAQP3, hsAQP7, hsAQP9, and hsAQP10. Besides exhibiting permeability for glycerol, certain aquaglyceroporins (e.g.,hsAQP3) allow hydrogen peroxide to pass and hence modulate intracellular H_2_O_2_ levels (Ma et al., 2023; Miller et al., 2010; Watanabe et al., 2016). In PfAQP, F138Y lies away from the predicted glycerol transport channel indicated by the five glycerol molecules in red stick format (**Figure 2E**), while in hsAQP3, the human equivalent residue (F161) lies near the H_2_O_2_ channel (Wragg et al., 2020) on TM4 and near the critical E loop (GFNSGY in human AQP3; GGNTGF in PfAQP) (**Figure 2F-H**). The closed conformation of human AQP3 results from loop E collapsing into the channel with Y212 extending to the NPA motifs at the channel core (Huang et al., 2025). This data suggests that the residue might be involved in the transition between open and closed conformations that occur when there is a change in pH or when H_2_O_2_ is present (Huang et al., 2025).

### CRISPR/Cas9 genome editing validates the mutation F138Y in aquaglyceroporin confers resistance to cryptosporin

To determine the independent contribution of the F138Y allele, CRISPR/Cas9 was used to introduce the F138Y into Dd2 parent as described previously (Adjalley & Lee, 2022). Successful editing was confirmed via PCR and Sanger sequencing. Upon cloning via limiting dilution, two clones, 2E2 and 2E3, were selected with the desired mutation. When phenotyped using a SYBR green I – based dose response assay, both clones had a 2-fold, significant EC50 shift from the parent (**Figure 2J** and **Table 3**), similar to the resistance shift observed in the *in vitro* evolved clones, while no significant EC_50_ shift was observed with the control compound atovaquone (**Figure 2K** and **Table 3**). The fact that F138Y alone confers resistance to cryptosporin is strong evidence that PfAQP mediates cryptosporin resistance. We also tested sensitivity of the edited lines to artemisinin. Not surprisingly, having no duplications of SOD-1 and SOD-2, 2E2F138Y and 2E3 F138Y did not show a resistance shift with artemisinin (**Figure 2L**, **Table 3**).

### Duplications of superoxide dismutase genes are observed in evolved clones

As part of our analysis, we also examined our evolved clones for structural alterations using a combination of increased read coverage and the presence of discordant paired-end reads. Increased read coverage is a standard method for identifying amplification CNVs in *P. falciparum*. Discordant paired-end reads which are observed when reads from near the boundaries of duplication event are mapped to a reference genome without the CNV, provide an orthogonal independent source of evidence for duplications in short read sequencing (Luth et al., 2024). In addition, discordant reads can be used to determine if the CNV is a tandem event as well as the exact location of the recombination event. Interestingly, all clones had at least one tandem duplication on chromosome 6 (**Table S5**). In addition, clones from flask No.2 (2-F7 and 2-F10) also had one additional duplication located on chromosome 8. Additionally, one clone (1B7) in flask No.1 also showed the chromosome 8 duplication with 32 RF (reverse-forward) discordant read pairs, suggesting the presence of a tandem duplication, despite minimal extra read coverage signal in the region (**Figure 3A**). The CNVs were of different sizes, suggesting that they all arose independently. Duplication of the chromosome 6 section and the chromosome 8 section were consistently observed across three independent bulk culture sequencing experiments prior to clone sequencing (**Table S5**; BioProject PRJNA1451291), suggesting that chromosome 6 amplification may arise earlier than chromosome 8 amplification, or that, low-frequency chromosome 8 events were not detected due to limited sequencing depth of bulk culture. Our database showed that these amplifications are highly specific to cryptosporin-evolved parasites and have not been previously detected in the thousands of whole genome sequences of compound-evolved parasites we have previously examined (Luth et al., 2024). No other structural alterations, including additional amplification of PfMDR1 or other drug resistance genes, were observed in the set.

**Figure 3.**
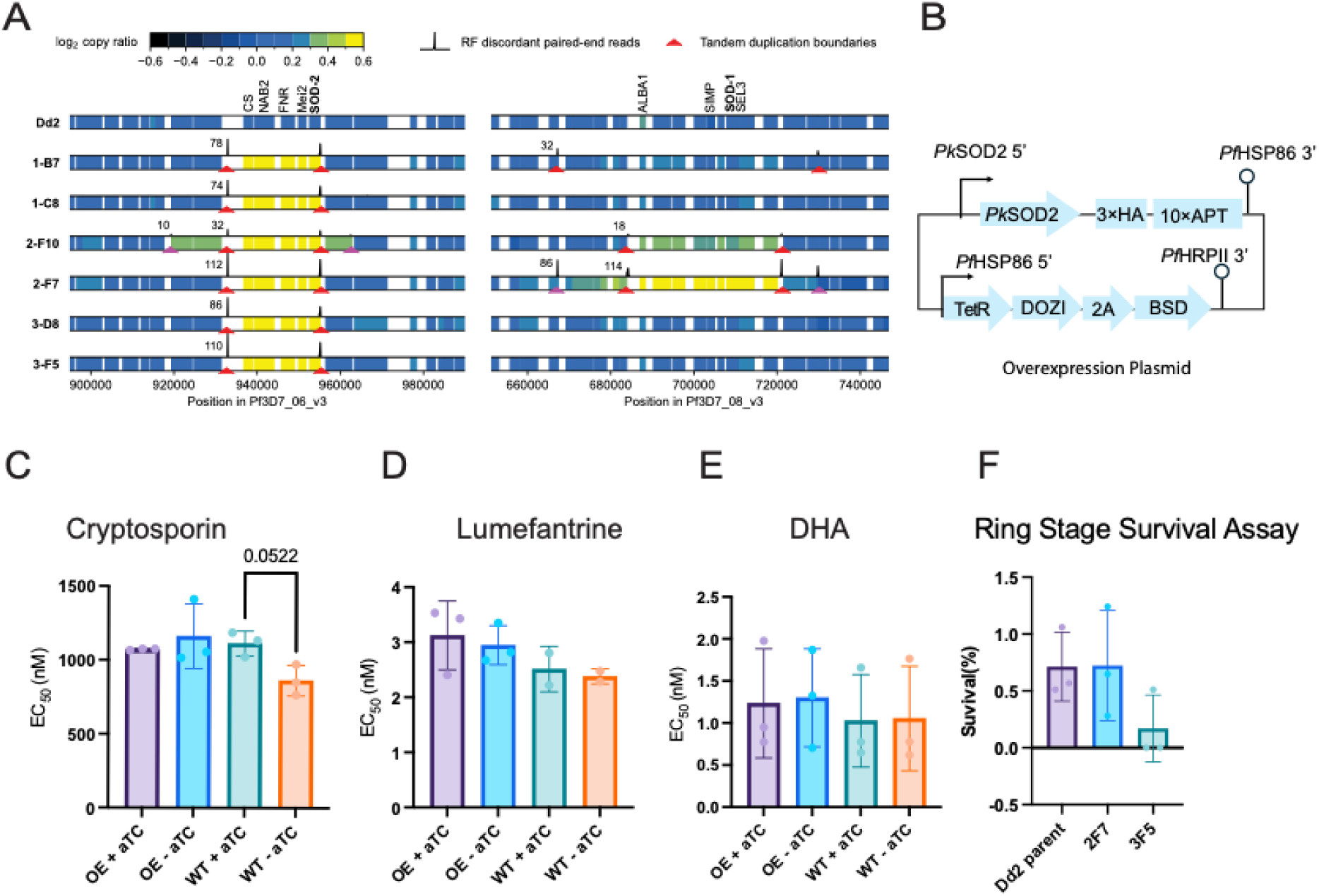
*In vitro* evolution of cryptosporin-resistant parasites reveals duplications of SOD-1 and SOD-2. (**A**) Coverage-based estimates of gene copy ratio and discordant read pair evidence for copy number variants (CNVs) from short read sequencing of the six cryptosporin-resistant clones and their Dd2 parent. CNVs were found in two genomic regions, one on chromosome 6 (left panel) and one on chromosome 8 (right panel). Gene blocks are colored according to log2 copy ratio. The number of reads belonging to reverse-forward (RF) discordant pairs is plotted as a black histogram (100 nt bin size). Tandem duplication boundaries are marked with a red or pink arrow below. The number labelled near the left peak in each pair is the total number of paired-end reads belonging to that pair. (**B**) Schematic of overexpression in *Plasmodium knowlesi*. (**C**) Effect of episomal PkSOD-2 overexpression on cryptosporin sensitivity in *Plasmodium knowlesi*. An episomal PkSOD-2 overexpression (OE) line was generated in *P. knowlesi* using a TetR-DOZI regulated expression system. Briefly, the coding sequence of PkSOD2 (lacking a stop codon) was fused in frame to a C-terminal 3 x HA tag and aptamer array. For transfection, 200 million Percoll purified schizonts were transfected with 10ug of plasmid using Amaxa 4D-Nucleofector (Lonza). Blasticidin-S HCl (5 µg/mL) was added 24h after transfection and maintained throughout the selection. Dose response assays were performed in the presence (+aTC) or absence (−aTC) of anhydrotetracycline (aTC) at 100nM in both the wild-type and the OE lines against cryptosporin (**C**) and the control compounds lumefantrine (**D**) and dihydroartemisinin (DHA) (**E**). Data represent mean ± SD from three biological replicates. (**F**) Ring stage survival assay showed that the in vitro evolved clones are not cross resistant to DHA. Two clones (2F7 and 3F5) with PfSOD-2 duplication, with or without PfSOD-1 duplication were selected for this assay. Briefly, parasites were synchronized with sorbitol over two consecutive intraerythrocytic cycles, followed by 75% Percoll purification of schizonts. Purified schizonts were allowed to invade for 3 h before treatment with DHA (700 nM) or DMSO (0.01%) for 6 h. Following treatment, cultures were washed three times to remove the drug and returned to standard culture conditions for an additional 66 h. Parasitemia was determined by Giemsa-stained thin blood smear and microscopy. At least 5,000 red blood cells were counted independently by three investigators for each treatment group. Data shown are mean ± SD of three biological replicates.

### Superoxide dismutase is common to both cryptosporin-induced CNVs

It can be challenging to identify the driver gene in CNVs, which typically have several candidates in the minimal region (here four to six in each). We thus searched for genes with missense mutations in the regions at allele frequencies of less than 1 but this did not reveal any candidates. Nevertheless, our analysis did show that both the chromosome 6 and chromosome 8 amplifications contained either mitochondrial superoxide dismutase (PfSOD-2, PF3D7_0623500), or cytosolic superoxide dismutase (PfSOD-1, PF3D7_0814900) (**Table 3**, **Figure 3A**). Superoxide dismutase is a critical antioxidant enzyme that protects cells by catalyzing the breakdown of harmful superoxide radicals into oxygen and hydrogen peroxide (Muller, 2004). The likelihood of this occurring by chance is very slim.

### Overexpression of PkSOD-2 in *Plasmodium knowlesi* does not confer a significant EC_50_ shift against cryptosporin

To investigate whether increased expression of SOD-2 modulates parasite susceptibility to cryptosporin, we utilized a conditional overexpression (OE) system in *P. knowlesi*. Overexpression of genes in *P. falciparum* can be challenging due to the AT-rich nature of *P. falciparum* genes. In addition, we found that cryptosporin produced an equivalent killing effect in *P. knowlesi*. In this system, episomal PfSOD-2 expression was regulated using the TetR-DOZI inducible platform controlled by anhydrotetracycline (aTC) ( **Figure 3B**). Drug sensitivity assays comparing wild -type and PkSOD2 OE parasites in the presence or absence of aTC showed only modest changes in cryptosporin sensitivity (**Figure 3C**). We also tested these conditions against lumefantrine, a control compound. PkSOD-2 OE parasites exhibited a slight increase in lumefantrine EC_50_ relative to wild-type parasites, however, the OE parasites without aTC also showed a slight increase in EC_50_ (**Figure 3D**). No significant change in sensitivity to DHA was observed with or without overexpression of PfSOD-2 (**Figure 3E**). These data suggests that overexpression of PkSOD-2 did not produce a significant EC_50_ shift against cryptosporin relative to wild-type parasites, suggesting that elevated PkSOD-2 expression alone is insufficient to confer strong resistance to cryptosporin. It’s well known that superoxide dismutase upregulation contributes to artemisinin resistance, we are interested to see whether the clones bearing SOD duplications have an altered ring stage survival rate. We conducted a ring stage survival assay. Among the two clones with PfSOD-2 duplication and with (2F7) or without (3F5) PfSOD-1 duplication, we did not observe apparent increased DHA survival rate in our (**Figure 3F**). Taken together, although we did not observed significant difference of compound sensitivity, SOD duplications may help with long term survival or tolerance and survival/tolerance may not be adequately captured in a 72-hour dose response assay.

### PfAQP permeability is unaffected by cryptosporin

To test if cryptosporin directly affects PfAQP permeability, we employed a strain of *Saccharomyces cerevisiae* in which the non-essential, native AQP (fps1) was deleted and replaced with the P. falciparum gene. We first assayed the growth of PfAQP-expressing BY4742 Δfps1 S. cerevisiae lacking the endogenous aquaglyceroporin Fps1 in methylamine/methyl ammonium media (Krenc et al., 2014). At a media pH of 5.6, the predominant protonated methyl ammonium form is taken up via the endogenous yeast ammonium transporters MEP1–3. Under the less acidic pH conditions of the yeast cytosol methyl ammonium partially dissociates into yeast-toxic neutral methylamine. Cell growth indicates methylamine efflux via PfAQP (Zeuthen et al., 2006), while blocking PfAQP inhibits this growth. Cells expressing pentamidine-sensitive TbAQP2 from Trypanosoma brucei (Song et al., 2016), and those lacking aquaglyceroporin expression served as controls.

Growth curves of PfAQP- (left) or the non-expressing cells (middle) with cryptosporin at varying concentrations (0, 0.01, 10 μM) were obtained (**Figure 4A**). TbAQP2-expressing cells (right) with or without pentamidine served as a control. Growth in methylamine media of TbAQP2-expressing cells was inhibited to the background level by adding 50 µM pentamidine, demonstrating suitability of the assay. Growth of PfAQP-expressing cells decreased at 10 µM cryptosporin.

**Figure 4:**
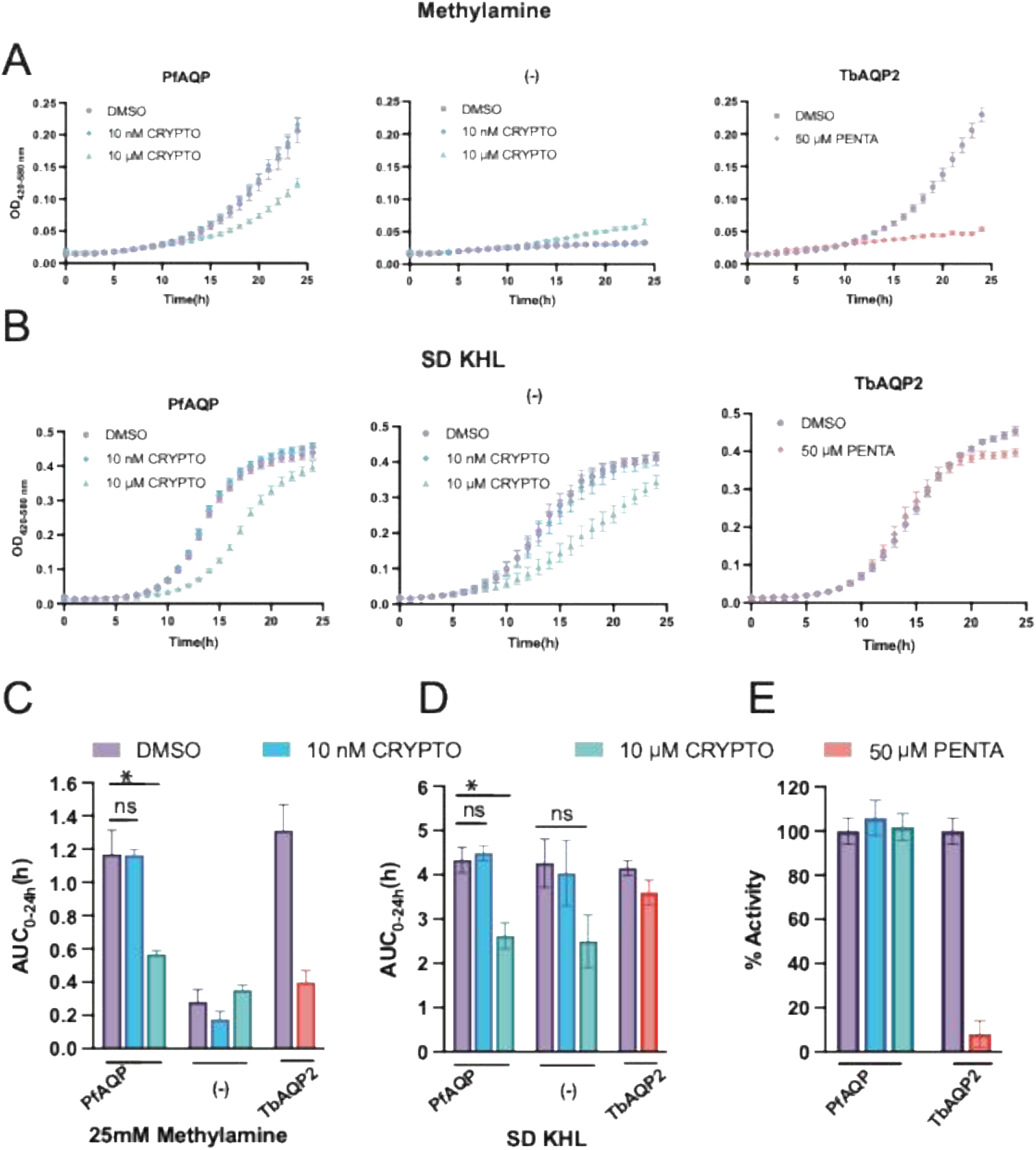
Assessment of the effect of cryptosporin on PfAQP permeability. **(A)** Phenotypic growth assays of BY4742 Δfps1 yeast expressing PfAQP or TbAQP2, and non-expressing cells (-). Growth curves were monitored in the presence(top) or absence (bottom) of methylamine, with and without cryptosporin or pentamidine as indicated. Growth under methylamine conditions indicates aquaglyceroporin functionality. Error bars denote SEM from 2 to 3 technical replicates of 3 biological replicates. **(B)** Quantification of the growth effects by the determination of the area under the curve (AUC). Statistical significance was assessed using unpaired t-tests (* p < 0.05). **(C)** Uptake of ^14^C-radiolabeled glycerol via PfAQP- or TbAQP2-expressing yeast in the absence or presence of cryptosporin or pentamidine. Uptake rates were measured over 1 min in a 1 mM inward glycerol gradient at pH 7.2, normalized to uninhibited conditions after subtraction of background uptake from non-expressing cells. Error bars indicate SEM from two biological replicates with two technical replicates.

However, the growth defect elicited by cryptosporin (10 µM) was evident in both PfAQP-expressing and non-expressing cells in nonselective SD KHL media without methylamine (Figure 4A). This indicates that the observed growth defect might not be due to the inhibition of PfAQP but rather the cytotoxicity of cryptosporin. For quantification, we determined the areas under the curve (AUC) over 24 h (**Figure 4B**). The AUC0-24 h of yeast expressing PfAQP was significantly decreased by about half in the presence of 10 µM cryptosporin in methylamine as well as in non-selective SD KHL media (p < 0.02), which is indicative of a general toxicity of cryptosporin in yeast rather than a direct effect on PfAQP.

Next, we employed a biophysical aquaglyceroporin permeability assay using 14C-radiolabeld glycerol (Geistlinger et al., 2022). Here, PfAQP- or TbAQP2-expressing Δfps1 yeast cells were subjected to a 1 mM inward glycerol gradient at pH 7.2. Background glycerol transmembrane diffusion of non-expressing cells (0.05 ± 0.01 nmol mg–1 min–1) was subtracted (**Figure 4C**). This yielded glycerol uptake rates of 0.60 ± 0.03 nmol mg–1 min–1 for PfAQP-expressing cells and 0.31 ± 0.01 nmol mg–1 min–1 for TbAQP2 in the absence of inhibitor (100% activity). Addition of 50 µM pentamidine (Song et al., 2016) almost fully inhibited TbAQP2-facilitated glycerol transport (8 ± 6 % activity), while glycerol permeation via PfAQP remained unaffected under treatment with 10 nM or 10 µM cryptosporin. Differing from the phenotypic assay, this rapid biophysical approach does not suffer from cytotoxic effects of putative inhibitor compounds. Taken together, the assays show that cryptosporin does not directly inhibit PfAQP-facilitated

## Discussion

*Plasmodium falciparum* is especially prone to oxidative stress due to the digestion of hemoglobin, during which, ferriprotoporphyrin IX(FP) and reactive oxygen species (ROS) are generated (Kavishe et al., 2017; Rosenthal & Ng, 2020). While most of the toxic FP is polymerized into crystalline hemozoin, some escape into the cytoplasm. Once there, FP can react with oxygen and form reactive oxygen species, which damages the cell membrane through oxidation of sulfhydryl groups and lipid peroxidation (Kavishe et al., 2017). It’s not uncommon for antimalarials to exert their activity by inducing oxidative stress. Quinoline drugs such as chloroquine and amodiaquine act through binding to FP and hindering its polymerization into hemozoin, leading to the accumulation of toxic FP (Fitch, 2004). In chloroquine-resistant parasites, elevated glutathione S-transferase (GST) activity has been reported (Deharo et al., 2003). Addition of antioxidant N-acetylcysteine, which is known to provide thiol groups and increase cellular glutathione level, desensitizes chloroquine in *Plasmodium berghei* (Deharo et al., 2003). Primaquine induces ROS in erythrocytes and leads to rapid depletion of glutathione and cytoskeleton damage, lipid peroxidation and hemolysis (Bowman et al., 2005). One of the major modes of action of artemisinin is through inducing oxidative stress. The levels of antioxidants and glutathione were shown to be reduced in artemisinin-treated parasite (Cui & Su, 2009). During the activation of artemisinin through cleavage of the endoperoxide bridge, free radicals and ROS were released. Artemisinin was found to form adducts with heme and FP which resulted in toxic FP accumulation and ROS production (Ma et al., 2021). In *P. berghei*, artemisinin was found to induce mitochondria ROS and superoxide dismutase activity in the mitochondria was elevated (Hou et al., 2020). Addition of N-acetylcysteine 2 h after exposure to artesunate resulted in abolition of artesunate activity in *P. falciparum* (Arreesrisom et al., 2007). Using transgenic lines with integration of redox sensors in mitochondria and cytosol, hydrogen peroxide level was shown to be elevated in parasites treated with artemisinin derivatives, quinine, and mefloquine in the mitochondria, and with chloroquine in the cytosol (Rahbari et al., 2017).

Most eukaryotic cells contain five major antioxidant enzymes - superoxide dismutase (SOD), catalase (CAT), glutathione peroxidase (GP), glutathione S- transferase (GST), peroxiredoxin (Jomova et al., 2023). *P. falciparum* lacks CAT and GP expression but expresses two SODs. Both SODs use iron as a cofactor with one found in cytosol (PfSOD-1, chromosome 8) and the other in mitochondria (PfSOD-2, chromosome 6) (Muller, 2004). Our RNA-seq data showed enrichment of genes associated with sexual development with short exposure to cryptosporin (1h), consistent with previous reports that oxidative stress induces gametocytogenesis in malaria parasites (Chaubey et al., 2014). Through the duplication of PfSOD-2 (all clones) or both SODs (clones 1B7, 2F10, and 2F7), *P. falciparum* may be able to counter the oxidative stress induced by cryptosporin. This highlights the role of SODs in antioxidant defense in these parasites. *In vitro* selected *P. falciparum* resistant to DHA showed upregulation of PfSOD-1 and PfSOD-2 along with elevated activities of constituents of antioxidant systems including glutathione and thioredoxin and GST (Cui et al., 2012). However, in our ring stage survival assay, clones (2F7 and 3F5) with PfSOD-2 duplication, with or without PfSOD-1 duplication, didn’t show an elevated DHA (700 nM for six hours) survival rate as compared to the parent Dd2. Cryptosporin likely induces significant oxidative stress in these parasites. The exact mechanism is unknown, however it is likely that iron can attack the oxygen in the tetrahydrofuran ring, resulting in the opening of the ring.

In addition to the duplications of SOD1 and SOD2, our study implicates PfAQP in the drug sensitivity of cryptosporin through resistant parasite generation. Introducing a single mutation (F138Y) in PfAQP via CRISPR/Cas9 rendered a near two-fold resistance to cryptosporin, highlighting the role of PfAQP in mediating cryptosporin sensitivity. This is the first report implicating PfAQP in drug resistance in *Plasmodium*. Unlike many other organisms that have multiple aquaporins, *P. falciparum* possesses one single aquaglyceroporin localized to the plasma membrane (Hansen et al., 2002). Oocyte swelling assay showed that PfAQP is highly permeable to water, glycerol, sugar alcohols up to five carbons, ammonia, and urea (Hansen et al., 2002). It is believed to facilitate glycerol uptake for lipid biogenesis and maintain osmotic balance (Beitz et al., 2004). PfAQP contains six full transmembrane helices and two half-helices that meet at the center of the pore. The two canonical NPA motifs capping the half-helices are replaced with NLA (aa. 70-72) and NPS (aa. 193-195), respectively (Newby et al., 2008)*. P. berghei* aquaglyceroporin (PbAQP) was found to be essential in both blood and liver stage. PbAQP-null blood stage parasites grew slower and were less infectious than wild-type parasites in addition to a deficiency in glycerol uptake into erythrocyte (Promeneur et al., 2007). PbAQP-null hepatic merozoites incorporated less glycerol into phosphatidylcholine and demonstrated slow growth (Promeneur et al., 2018). Aquaglyceroporins in other protozoa parasites such as *Trypanosoma* mediate drug sensitivity. *Trypanosoma brucei* TbAQP2 was shown to directly facilitate the uptake of pentamidine and melarsoprol (Baker et al., 2012). High-affinity binding of pentamidine to TbAQP2 has been demonstrated experimentally (Song et al., 2016) and more recently confirmed through cryo-electron microscopy structural studies (Chen et al., 2024; Matusevicius et al., 2026). The ability of TbAQP2 to bind drug molecules is attributed to the nonconventional NSA/NPS motifs and an unusual, larger selectivity filter with leucine replacing arginine at residue 264(Chen et al., 2024). The L264R mutation abolished the dicationic pentamidine binding, leading to drug resistance (Alghamdi et al., 2020).

One of the main roles of PfAQP is glycerol uptake from host serum into erythrocytes, which is important for parasite virulence. Knocking out of human erythrocytic AQP9, a major pathway for glycerol uptake, increased the resistance of mice to *P. berghei* (Liu et al., 2007). If cryptosporin hinders PfAQP glycerol permeability, parasites could shift to glycolysis for glycerol-3-phosphate required for glycerolipid synthesis (Hansen et al., 2002). The heavy reliance on glycolysis could generate oxidative stress (Atochina-Vasserman et al., 2013). Recent studies have hypothesized that glycerol production is an antioxidant defense mechanism used by various organisms to maintain redox homeostasis. Under anaerobic conditions, the production of glycerol was observed in protozoan parasites such as *Trypanosoma* and *Leishmania* but not human erythrocytes (Darling et al., 1988). If cryptosporin does hinder glycerol flux, it could cause oxidative stress either via elevated glycolysis or by hindering glycerol-mediated detoxification. However, our permeability assay showed that the glycerol uptake by PfAQP was unaffected by cryptosporin, suggesting that PfAQP mediates cryptosporin sensitivity via a different mechanism.

Members of the AQP3 ortholog group mediate the transmembrane diffusion of H_2_O_2_, contributing to anti-oxidative stress response and signaling transduction (Bienert & Chaumont, 2014). PfAQP facilitates the transport of H_2_O_2_ albeit with relatively low conductance (Almasalmeh et al., 2014). Recently, single particle cryo-EM structures of human AQP3 showed that H_2_O_2_ regulates AQP3 channel closure independent of pH(Huang et al., 2025). At pH=8, excess H_2_O_2_ triggered AQP3 closure with extracellular loop E (GFNSGY, residue 207-212) collapsing into the pore with Y212’s hydroxyl group extending to NPA motifs. The hydrogen bonding network of hydrogen peroxide stabilized the closed conformation of AQP3 which places AQP3 as a redox sensor and regulator. F138 in PfAQP is not directly lining the H_2_O_2_ channel but is rather located in the extracellular vestibule facing the membrane side. However, its location was found to be close to N140 (corresponding to D163 in human AQP3), S131(corresponding to H154 in human AQP3) and loop E (GGNTGF) which suggests that it could play a role in stabilizing a certain conformation of PfAQP. In human AQP3, protonation of D163 triggers a cascade of conformational changes, beginning with disruption of the D163–N209 interaction and subsequently the HFHF stacking network (H154–F208–H53–F56), ultimately causing loop E to collapse into the channel pore and block the channel. We hypothesize that the mutation F138Y (from a hydrophobic phenylalanine to partial hydrophilic tyrosine and with an additional hydroxyl group) renders a conformational change in PfAQP which affects the closing and opening of the pore allowing hydrogen peroxide permeability, likely contributing to a resistance mechanism.

## METHODS

### Fungal isolate and fermentation

*Acaromyces ingoldii* (internal code USFS 08041501) was isolated from a loblolly pine bolt from a tree infected with bark beetles in the Kisatchie National Forest in Louisiana, US (Olatinwo & Fraedrich, 2019; Olatinwo et al., 2019). The isolate was previously identified using ITS sequencing (Olatinwo & Fraedrich, 2019; Olatinwo et al., 2019). For chemical studies, the fungus was cultured in mycobags (Unicorn Bags, TX US) on a monolayer of Cheerios supplemented with 0.3% sucrose and 0.05% chloramphenicol for 4 weeks.

### *Plasmodium* Blood Stage Culture

Parasites in the Winzeler and Chakrabarti labs were grown based on protocol by Trager and Jensen with slight variations per lab (Trager & Jensen, 1976). The Chakrabarti lab utilized RPMI 1640 with 25 mM HEPES pH 7.4, 26 mM NaHCO_3_, 15 mg/L hypoxanthine, 0.5% Albumax II, 25 mg/L gentamycin, and 2% dextrose. Parasites were grown in human A+ erythrocytes at 37°C and 5% CO_2_, 5% O_2_, 90% N_2_, or 95% air. The Winzeler lab utilized RPMI 1640 with 26 mM NaHCO_3_, 0.1 mM hypoxanthine, 0.25% Albumax I, and 50 µg/L gentamicin. Parasites were grown in human O+ erythrocytes at 37°C with 5% CO_2_, 3% O_2_, and 92% N_2_.

### Blood Stage Screening

Preliminary screening of the natural product library including cryptosporin was accomplished using a SYBR Green I-based assay performed by the Chakrabarti lab. Protocol was based on established procedure by Smilkstein et al (Smilkstein et al., 2004) as described in a previous publication(Lee et al., 2021). Briefly, asynchronous blood stage parasites were plated at 1% parasitemia, 1% hematocrit and incubated with compound for 72 h. Plates were then frozen and thawed to promote lysis prior to the addition of 1x SYBR Green I in a buffer of 20 mM Tris-HCl, 0.08% saponin, 5 mM EDTA, and 0.8% Triton X-100. After a 45-60 min incubation at room temperature, florescence at 485 nm excitation, 530 nm emission was determined with a Synergy Neo2 plate reader (BioTek Winsooki, VT). Screening in ScDHODH line performed by the Winzeler lab as per previous publication(Collins et al., 2023). In brief, asynchronous Dd2-ScDHODH(Painter et al., 2007), a transgenic line expressing *S. cerevisiae* cytosolic type I DHODH (**Table S6**), was plated at 0.3% parasitemia, 2.5% hematocrit. Parasites were grown with or without attB, and with or without attB with proguanil (PG). Compound was added using a MultiFlo™ microplate dispenser (BioTek Winsooki, VT) and plates were incubated for 72 h. At this time, a solution of 10 x SYBR Green I in a buffer of 20 mM Tris/HCl, 5 mM EDTA, 0.16% (w/v) saponin, and 1.6% (v/v) Triton X-100 was added to each well using the MultiFlo™ dispenser. After incubation at room temperature for 24 h, fluorescence was read at 485 nM excitation, 530 nm emission using a EnVision® Multilabel Reader (PerkinElmer Waltham, MA). Dose response curves for both labs were generated using CDD Vault (https://www.collaborativedrug.com/).

### Liver Stage Screening

Screening against *Plasmodium* liver stage was performed based on established protocols (Antonova-Koch et al., 2018; Collins et al., 2023). In brief, human HepG2-A16-CD81 cells were infected with fresh *P. berghei* sporozoites (*P. berghei* ANKA GFP-Luc-SMcon, **Table S6**) following dissection of *A. stephensi* salivary glands. Compound was incubated with HepG2 cells at 37°C, 24 h pre-invasion. At this time, sporozoites were added at a density of 1 × 10^3^ cells per well, and plates were incubated for an additional 48 h. Following this, luciferin reagent (Promega BrightGlo) was added for luciferase detection using a EnVision® Multilabel Reader (PerkinElmer Waltham, MA). For comparison, HepG2 cells were seeded with compound as described above without the addition of sporozoites. After 72 h, cytotoxicity was determined using a Promega CellTiterGlo^®^ luminescence assay. Dose response curves were generated using CDD Vault (https://www.collaborativedrug.com/).

### Minimum Inoculum of Resistance Determination

MIR determination was performed using synchronous Dd2-B2 clonal parasites at 0.3% parasitemia, 1% hematocrit. Cryptosporin or DSM265 control were added at 3 × EC_90_ to ring-stage parasites with inocula of 1.4 × 10^7^ (12 wells) or 3 × 10^7^ (3 flasks). Cultures were monitored for 60 days for parasite recrudescence using Giemsa blood smears and flow cytometry with SYBR Green and MitoTracker Red on an iQue flow cytometer (Sartorius, Göttingen, Germany).

### Stage Specific Assay

Assay was performed on synchronous Dd2 culture as previously described (Collins et al., 2021). Parasite synchronization was achieved through combination of MACS column magnetic separation (Mata-Cantero et al., 2014) and sorbitol synchronization (Lambros & Vanderberg, 1979). Once synchronous, cultures were exposed to a 5 × EC_50_ concentration of cryptosporin or a vehicle control at either 6, 18, 30, or 42 HPI. Afterwards, samples were collected at 12 h intervals until 54 HPI for fixing in paraformaldehyde and staining via Giemsa. Fixed samples were then stained for flow cytometry with YOYO-1 and read on a CytoFLEX s (Beckman Coulter, Brea, CA) at 488 nm excitation with 500,000 events per well. Data was then analyzed with FlowJo version 10 using uninfected RBCs and unstained infected RBCs as gating controls. Median and total fluorescence values were extracted and plotted in Graph Pad Prism (version 10) for each replicate with mean and SEM. Two biological replicates were performed in total. Significance between the vehicle control and treated wells for each timepoint was determined using a multiple comparison test with a two-way ANOVA. As there was only one timepoint for the 42 HPI treatment (collection at 54 HPI), a paired t-test compared to the 54 HPI control was used to determine significance for this point.

### Rate of Killing Assay

The killing rate assay was performed as previously described (Collins et al., 2021) in semi-synchronous Dd2 culture in the majority ring stage. Cryptosporin, fast acting control DHA, or vehicle was added to at 10 x EC_50_ to culture at 1% parasitemia, 4% hematocrit. Compound or control was incubated for 12, 24, or 48 h prior to compound wash off with RPMI. Parasites were then monitored at wash off and every 24 h after for a total of 5 days. Samples were taken every 24 h for Giemsa stain and flow cytometric analysis with SYBR Green I and MitoTracker Deep Red FM using a CytoFLEX S (Beckman Coulter, Brea, CA) with 100,000 events per well. Data was then analyzed on FlowJo version 10 with a no MitoTracker control, uninfected RBC control, and no staining control for gating. Populations identified as both “SYBR positive” and “MitoTracker positive” were used to track the percentage of viable parasites. Three biological replicates were then graphed using Graph Pad Prism( version 10) with error bars representing mean and SEM across replicates.

### RNA Extraction

RNA was collected for RNA-Seq and RT-qPCR as previously described (Collins et al., 2023) from synchronous, 3D7 culture at 10% parasitemia, 4% hematocrit. Parasites were incubated with 1 × EC_50_ concentration of inhibitor in late trophozoite prior to saponin lysis and 2×PBS wash. Pelleted parasites were then resuspended in TRIzol^®^ reagent and RNA was extracted using Direct-zol™ RNA Miniprep Plus (Zymo Research Corporation, Irvine, CA.) according to manufacturer’s protocol. RNA concentration and integrity was then confirmed via nanodrop and gel electrophoresis prior to integrity checks with Agilent Technologies 2100 Bioanalyzer.

### RNA-Seq

After RNA integrity confirmation, ribosomal RNA depletion, and RNA fragmentation, library preparation was performed per Illumina TruSeq-stranded-total-RNA-sample preparation protocol. Paired-end sequencing was achieved with Illumina’s NovaSeq 6000 sequencing system with Agilent Technologies 2100 Bioanalyzer High Sensitivity DNA Chip for quality control. Cutadapt (Martin, 2011) was used to filter low quality reads during the transcript assembly process, in addition to LC sciences in house perl scripts. At this step, FastQC (Andrews, 2010) was used to confirm sequencing quality. Bowtie2 (Langmead & Salzberg, 2012) and HISAT2 (Kim et al., 2015) were utilized to map transcripts to the *P. falciparum* genome, with StringTie (Pertea et al., 2015), perl scripts, and gffcompare (Pertea & Pertea, 2020) used for assembly. Transcript expression or fragments per kilobase per million (FPKM) were determined as:

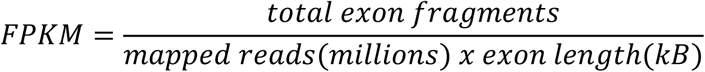

To determine significance, a parametric F-test comparison of nested linear models was performed with EdgeR(Robinson et al., 2010). Differential expression of transcripts was performed by considering each isoform individually (transcript level) or all transcripts of a single gene together (gene level).

### RT-qPCR

RNA extracted after cryptosporin incubation underwent cDNA synthesis with 100 ng of random hexamers following the manufacturers protocol for SuperScript First-Strand Synthesis System (Invitrogen, Waltham, MA). After, RT-qPCR was performed following manufacturers protocol for Select Master Mix (Life Technologies, Carlsbad, CA) with 400 nM of forward and reverse primers. Primers can be found in Table S7.

### Kinetic Solubility

Kinetic solubility testing was performed as previously described (Collins et al., 2021) using HPLC-UV. Compound was diluted to a final concentration of 200 µM in 2% DMSO and 1 × PBS, pH = 7.4 and added to Millipore solubility plates with 0.45 µM polycarbonate filter membranes. Plates were sealed and incubated for 24 h at room temperature, prior to vacuum filtration and filtrate collection. HPLC analysis was performed on an Agilent 1290 (Santa Clara, CA) and quantified using a calibration curve and linear regression in Agilent OpenLab Intelligent Reporting Software.

### Microsomal Stability

Microsomal stability testing of cryptosporin was performed as previously described (Collins et al., 2021). In brief, compound was tested in mouse microsomes from Menoxtech Male (CD-1), LN:1710069 mice. A solution of 0.63 mg/mL protein, 100 mM KPO_4_, and 1.3 mM EDTA was incubated with 0.1 mM compound for T = 0 minutes, T = 60 minutes, or T = 60 min without NADPH. Reactions at T = 0 were quenched using cold MeOH with a NADPH regenerative solution. Reactions at T = 60 had NADPH added to the reaction prior to incubation for 1 h followed by quenching with cold MeOH. For T = 60 without NADPH, water was added in place of NADPH. Plates were then centrifuged at 3000 x RPM at 4°C, and the percentage of compound remaining was determined using LC-TOFMS.

### Pharmacokinetics

To determine the pharmacokinetic parameters of cryptosporin, compound was dissolved in 100 µL of PBS (pH = 7.4) and delivered via intraperitoneal (IP) injection at a concentration of 20 mg/kg to four female BALB/c mice. Blood was collected by retro orbital bleeding at 0.5, 1, 2, 4, 8, and 24 hours post injection for analyte peak counts to determine the blood concentration. Values were then transformed by a non-compartmental analysis in RStudio (http://www.rstudio.com/).

### *In vitro* evolution of resistance in *P. falciparum*

The *in vitro* evolution experiments were conducted using *Plasmodium falciparum* Dd2 line. Parasites were cultured in human O-positive blood cells with leukocyte depleted (BioIVT, NY). Parasites were cultured in T75 flasks with 40 ml of media at 2% hematocrit, >3% parasitemia with or without cryptosporin. A single parental line (without drug addition, control) and 3 drug-treated parasite lines (3 biological replicates) were maintained at any given time. Every 2 or 3 days, parasitemia was determined via Giemsa smear and reset (if higher than 2 or 3%) to 2% (for a 3-day split, 1.6 ×10^8^ parasites) or 3% (for a 2-day split, 2.4 ×10^8^). If the parasitemia was not higher than 0.5%, drug was taken off, and parasites were allowed to recover. Parasites were replenished with fresh blood and complete RPMI 1640 media and appropriate amount of cryptosporin was added. Parasites were incubated at 37°C in sealed flasks filled with a mixture of gases containing 5% CO_2_, 3% O_2_, and 92% N_2_. In this manner, parasites were cultured continuously with increasing drug concentration from 40 nM to 2 μM for 10 months. Resistance development was monitored every 3 months through SYBR green I- based EC_50_ dose response assay as described above.

### Whole genome sequencing analysis

Genomic DNA was extracted from resistant clones obtained through limiting dilution as well as the Dd2 parent line. DNA libraries were generated using Nextera XT DNA Library Preparation Kit (Illumina), pooled, and sequenced on Illumina NovaSeq X Plus 10B flow cell in the 100 bp paired-end read (PE100) configuration. Reads were aligned to *Plasmodium falciparum* reference genome 3D7 (PlasmoDB v13.0) using an in-house pipeline (Cowell & Winzeler, 2019). SNVs and indel variants were called using GATK 4.0 HaplotypeCaller; confident mutations in resistant clones were identified by comparing parent and offspring allele depths using Fisher’s exact test (*P* < 0.001). To assess CNVs, genic coverage was standardized and denoised against a Dd2 panel of normals (constructed from 30 non-drug-selected Dd2 parents) using GATK DenoiseReadCounts. Contiguous gene intervals with log2 denoised copy ratio of >0.6 were considered to be amplified, while those with copy ratio < -0.6 were considered to have deamplification (copy loss).

### Limiting dilution to generate clonal lines

Parasites were diluted to 0.5 parasite per well into 96-well plates. Parasites were allowed to grow to detectable concentration over the next three weeks. Every 2 or 3 days, fresh media and blood was added to each well. Parasitemia was monitored starting from two weeks via Giemsa staining of thin blood smear every 2 or 3 days until viable parasites were identified. Two clones from each flask were selected. The resistance phenotype of these clones was measured via SYBR green I- based EC_50_ dose response assay as described below. DNA of the clones was extracted, and the library was prepped and sent for sequencing using the method described above. Sequencing analysis was done using the Winzeler lab in-house pipeline as described earlier.

### EC_50_ dose response assay in *P. falciparum* and *P. knowlesi*

The EC_50_ assay was performed at 1% parasitemia, 1% hematocrit with SYBR green I in 96-well plates. Briefly, the compound was subjected to a 3-fold serial dilution with ten dilution points. Three technical replicates were set up at each dilution point. The Dd2 parent line and the resistant parasites at 2% hematocrit and 1% parasitemia were added to the drug media at 1:1 ratio. The plate was incubated at 37°C for 72 hours in a chamber filled with a hypoxic gas mixture (3% oxygen, 5% carbon dioxide, and 92% nitrogen). After incubation, the cells were lysed overnight with lysis buffer (20 mM Tris-HCL, 0.08% saponin, 5 mM EDTA, and 0.8% Triton X-100) containing SYBR green I (10,000×, Thermo Fisher, cat # S7563). Fluorescence was measured at 485nm excitation and 530nm emission using a PHERAstar FSX plate reader (BMG Labtech, Germany). Dose response curves were graphed with Prism 9 (Version 9.5.1).

EC_50_ dose response assay using *P. knowlesi* yH1 parasites was performed as described above with the following differences. The drugs were printed directly onto 96-well black clear-bottom plates (Corning) using an HP D300e digital dispenser to generate a 2-fold serial dilution series, after which parasites were added at 1% hematocrit and 0.5% parasitemia in a final culture volume of 100 μL per well. Cultures were incubated at 37°C for 48 hours under the same gas conditions. Following lysis in SYBR green I-containing buffer as described above, fluorescence was measured using a SpectraMax iD5 microplate reader (Molecular Devices) at 494 nm excitation and 530 nm emission, and EC_50_ values were calculated in the same fashion as described above.

### CRISPR/Cas9 editing of PfAQP (PF3D7_1132800)

F138Y was introduced into the parental Dd2 line using methods previously described (Adjalley & Lee, 2022). The CRISPR guide 5’- GCTCCAGTTAAACTTATGGA was ligated into the BbsI site of the plasmid pDC2-coCas9-gRNA and verified by Sanger sequencing. A donor template with homology to the target site containing the F138Y point mutation (ttt → tat) was synthesized (Thermo Fisher) and ligated into the AatII and EcoRI sites of the same plasmid. Once constructed, this all-in-one plasmid should contain four critical components, namely, a SpCas9 codon-optimized for *P. falciparum*, gRNA expression cassette driven by U6 promoter, a donor template, and a drug selection marker (h*dhfr*). In addition, it also contains an ampicillin resistant marker for bacteria propagation. Transfections of Dd2 were performed as previously described (Fidock & Wellems, 1997). Briefly, sorbitol synchronized ring stage parasites at >5% parasitemia, 3% hematocrit were mixed with 50µg of plasmid in Cytomix (120 mM KCl, 0.15 mM CaCl_2_, 10 mM KH_2_PO_4_, 25 mM HEPES, 2 mM EGTA, 5 mM MgCl_2_, pH = 7.6) and transferred to a cuvette (0.2 cm gap) (Bio-Rad) and electroporated with Gene Pulser electroporator with Capacitance Extender (Bio-Rad) at 0.310kV and 950µF. Drug selection with 5 nM WR99210 (MedChemExpress, CAS No. 47326-86-3) was carried out for 10 days until no viable parasites were observed. Culture was maintained until parasite recrudescence. Returning parasites were confirmed for integration of donor template by PCR (**Table S7**) and Sanger sequencing before undergoing cloning by limiting dilution. Two clones were selected and genotyped by PCR and Sanger Sequencing. Once confirmed to have the F138Y mutation, the two clones were phenotyped along with parent Dd2 using SYBR green I – based dose response assay as described above.

### Plasmids and yeast transformation

Expression constructs of PfAQP in pDR196 and TbAQP2 in pRS426met25 were described previously (Song et al., 2016; Zeuthen et al., 2006). *Saccharomyces cerevisiae* BY4742 Δfps1 yeast cells lacking the endogenous aquaglyceroporin Fps1 were transformed using the lithium actetate/single-stranded carrier DNA/polyethylene glycol method (Gietz et al., 1995). Cells were grown in synthetic defined (SD) media containing 2 % (*w/V*) glucose, supplemented with 5 mg ml^−1^ ammonium sulfate, 20 mg ml^−1^ lysine-HCl, 20 mg ml^−1^ histidine and 100 mg ml^−1^ leucine (KHL), pH 5.6.

### Phenotypic yeast growth assay in methylamine media

BY4742 Δfps1 cells carrying PfAQP, TbAQP2 or empty vector DNA were grown in SD KHL media. Cells were harvested at 4000 *g*, 4 °C for 5 min, and diluted to a starting OD_600_ of 0.01 in SD KHL with or without 25 mM methylamine (plus 1 mg ml^−1^ proline replacing ammonium sulfate as nitrogen source). Cells were placed in 100-well plates (Honeycomb, Oy Growth Curves Ab Ltd.) with cryptosporin (10 nM or 10 µM) or pentamidine (50 µM) from a DMSO stock (1% final concentration). Growth at 29 °C was monitored over 24 h with changes in turbidity in the range of 420–580 nm (Bioscreen C, Oy Growth Curves Ab Ltd., with 10 s shaking before measurements at medium intensity). Measurements were done in 2–3 biological replicates with technical triplicates. The absorbance of cell-free medium was subtracted afterwards and the areas under the curves (AUC) were calculated by trapezoidal rule. Means differing more than 3 standard deviations were deemed outliners (Graf-Henning’s test for outliers) and thus excluded.

### Uptake of ^14^C-glycerol

BY4742 Δfps1 yeast expressing PfAQP or TbAQP2 were assayed for uptake of glycerol spiked with ^14^C-labeled glycerol (Geistlinger et al., 2022). Cells were grown in SD KHL to an OD_600_ of 0.9–1.4, harvested at 4000 *g*, 4 °C for 5 min, washed once with ice-cold 50 mM HEPES/Tris buffer (pH 7.2), adjusted to an OD_600_ of 50 ± 5 in this buffer, and kept on ice. Eighty-eight microliters of cell suspension were preincubated with cryptosporin (final concentration: 10 nM or 10 µM) from a DMSO stock (final concentration: 1%) for 30 min, or with pentamidine (final concentration: 50 µM) for 10 min. Following a 2-minute prewarming to 19 ± 1 °C, uptake was initiated by adding 22 µl of a 5 mM glycerol solution containing ^14^C-glycerol (Hartmann Analytic), creating a 1 mM inward glycerol gradient with a radioactivity spike of 0.002 µCi µl–1. After 1 min, 100 µl of the cell suspension were vacuum filtrated (0.45 µm GF/C filter membrane, Whatman) and washed with 7 ml of ice-cold water. The filters were transferred into vials with scintillation cocktail (Rotiszint eco plus, Carl Roth) and analyzed (Packard TriCarb 2900 TR, PerkinElmer Inc.). The assay was done in two biological replicates with technical duplicates. Background glycerol diffusion was determined using non-expressing cells and subtracted. Activity was normalized to uninhibited uptake of glycerol.

### Ring Stage Survival Assay (RSA)

The Dd2 parental line and selected clones (2F7 and 3F5) generated through *in vitro* evolution were subjected to a ring stage survival assay (Witkowski et al., 2013). Briefly, parasites were synchronized with 5% sorbitol over two consecutive cycles. Thirty hours later, schizonts were purified using a 75% Percoll gradient prepared with 15 volumes of heparinized RPMI (7.5 U/mL heparin) and 75 volumes of 90% Percoll. Parasitized blood was overlaid on top of 75% Percoll and spun at 1000 × g for 15mins. The interphase layer containing schizonts was collected and washed three times with RPMI medium. Purified schizonts were transferred to a T75 flask containing fresh erythrocytes at 2% hematocrit and incubated for 3 hours to allow invasion. Parasitemia was determined by Giemsa-stained thin smear microscopy. Cultures were then adjusted to 1% parasitemia and 2% hematocrit in a 24-well plate. Drug-treated wells received 700 nM (DHA, while control wells received 0.007% DMSO. After 6 h, DHA and DMSO were washed off, and parasites were cultured for an additional 66 h. Final parasitemia was determined by Giemsa-stained thin smears. Survival rate was calculated as the ratio of parasitemia in DHA-treated cultures relative to non-treated controls. Graphs were generated using Prism 9 Version 9.5.1.

### Generation of episomal PkSOD-2 overexpression line

An episomal overexpression plasmid was generated by Gibson assembly of the PCR-amplified PkSOD2 (PKNH_1126700) coding sequence together with 1 kb of upstream sequence (used as the promoter) into AatII- and AscI-digested pKD plasmid (Rajaram et al., 2020). The PCR fragment was amplified using the primers (**Table S7**). In the resulting construct, the PkSOD2 coding sequence (lacking a stop codon) is fused in frame to a C-terminal 3 x HA tag and TetR-regulated aptamer array. The pKD backbone also encodes TetR-DOZI and a blasticidin deaminase (BSD) resistance cassette. The plasmid was introduced into parasites using an Amaxa 4D-Nucleofector (Lonza) with the P3 Primary Cell 4D-Nucleofector X Kit L (Lonza, V4XP-3024), as previously described (Moon et al., 2013). Briefly, 200 million schizonts, purified on a 60% Percoll gradient (800 ×g for 20 min), were mixed with 10 µg plasmid DNA in 100 µL complete P3 primary cell solution and electroporated using program FP158. Blasticidin-S HCl (5 µg/mL) was added 24 hours after transfection and maintained continuously for selection. The sensitivity of cryptosporin, lumefantrine, DHA in wild-type and overexpression lines with or without anhydrotetracycline (aTC, 100nM) was determined via dose response assays (EC_50_ assays) as described above.

### Author Contributions

E.A.W., D.C., R.H.C. designed the research. T.J., J.E.C., S.B., B.T., R.C.S.E., L.T.F., C.L., N.M., R.P., N.M.S., J.D.M. performed the research. J.W.L., K.W., E.B., M.L., M.T.D., D.A.F, contributed to new reagents or analytic tools. D.W.C., T.J., J.E.C., J.B., S.B., analyzed the data. T.J., J.E.C., E.A.W. wrote the paper.

## Acknowledgments

This work was supported by grants from NIH R01 AI154777 (D.C., R.H.C., and E.A.W.), R21 AI143052 (D.C. and R.H.C.), and the Bill & Melinda Gates Foundation in support of the Malaria Drug Accelerator (MalDA) to E.A.W. and J.N. (OPP1054480). The PfAQP permeability assay conducted by S. B. was supported by Dr.Hilmer Stiftung grant (T0146 – 42.804). This publication includes data generated at the UC San Diego IGM Genomics Center utilizing an Illumina NovaSeq 6000 that was purchased with funding from a National Institutes of Health SIG grant (#S10 OD026929). The work conducted by B.T. was supported by grant (P500PB_225631) from the Swiss National Science Foundation. In addition, the authors would like to thank Dr. Rabiu Olatinwo for providing fungal samples to Dr. Robert H. Cichewicz’s lab.

## Data Availability

Whole genome sequencing data can be found online in SRA (Sequence Read Archive), BioProject PRJNA1451291. The dataset includes whole-genome sequencing at different timepoints (5, 7, 8, and 10 months) following resistance selection, including bulk-culture sequencing results (5, 7, and 8 months post-selection) and clone sequencing data (10 months post-selection).

