## supplemental materials for "Duplication of superoxide dismutase and a mutation in aquaglyceroporin mediates the sensitivity of *Plasmodium falciparum* to cryptosporin, a natural product derived from *Acaromyces ingoldii*"

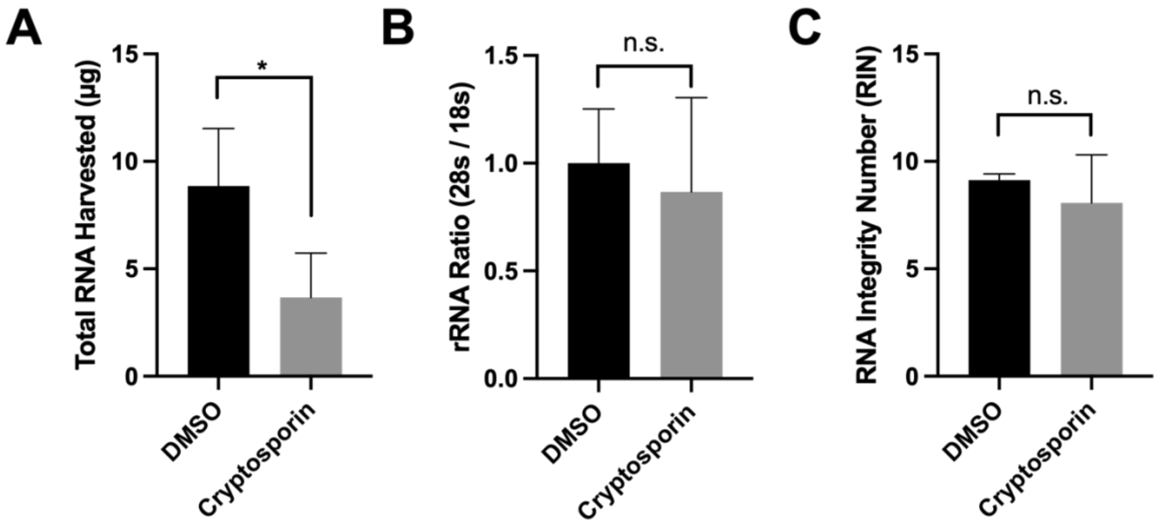

**Figure S1. RNA extraction following 1 h incubation with DMSO or cryptosporin.** Synchronous 3D7 culture was treated in trophozoite for 1 h with 5% DMSO or an $EC_{50}$ concentration of cryptosporin. **(A)** Total RNA collected from 30 mL of culture at ~10% parasitemia following incubation. **(B)** Ribosomal RNA ratio (28s/18s) of RNA following incubation. **(C)** RNA integrity number (RIN) following incubation. Data represents the mean and SEM of three replicates.

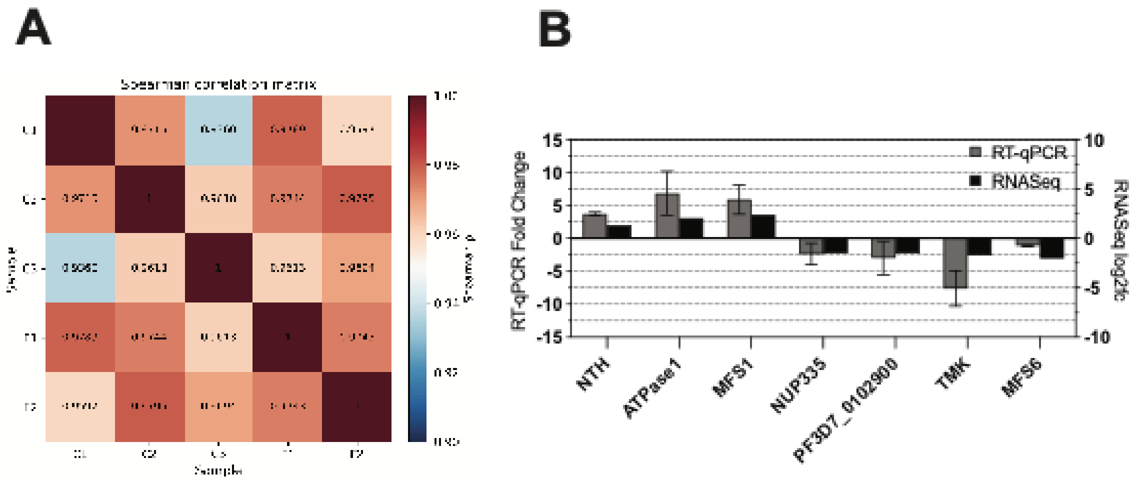

**Figure S2. Sample-to-sample reproducibility and RT-qPCR validation of RNA-seq results.** (A)Spearman correlation matrix of the five samples, namely, DMSO controls(C1-3) and cryptosporin-treated samples(F1-2). (B) RT-qPCR validation of differentially expressed transcripts compared to RNAseq log2fc. RT-qPCR data represents the mean and SEM of 2 biological replicates using housekeeping genes serine tRNA ligase (PF3D7_0717700) and 60S ribosomal protein L18-2 (PF3D7_1341300).

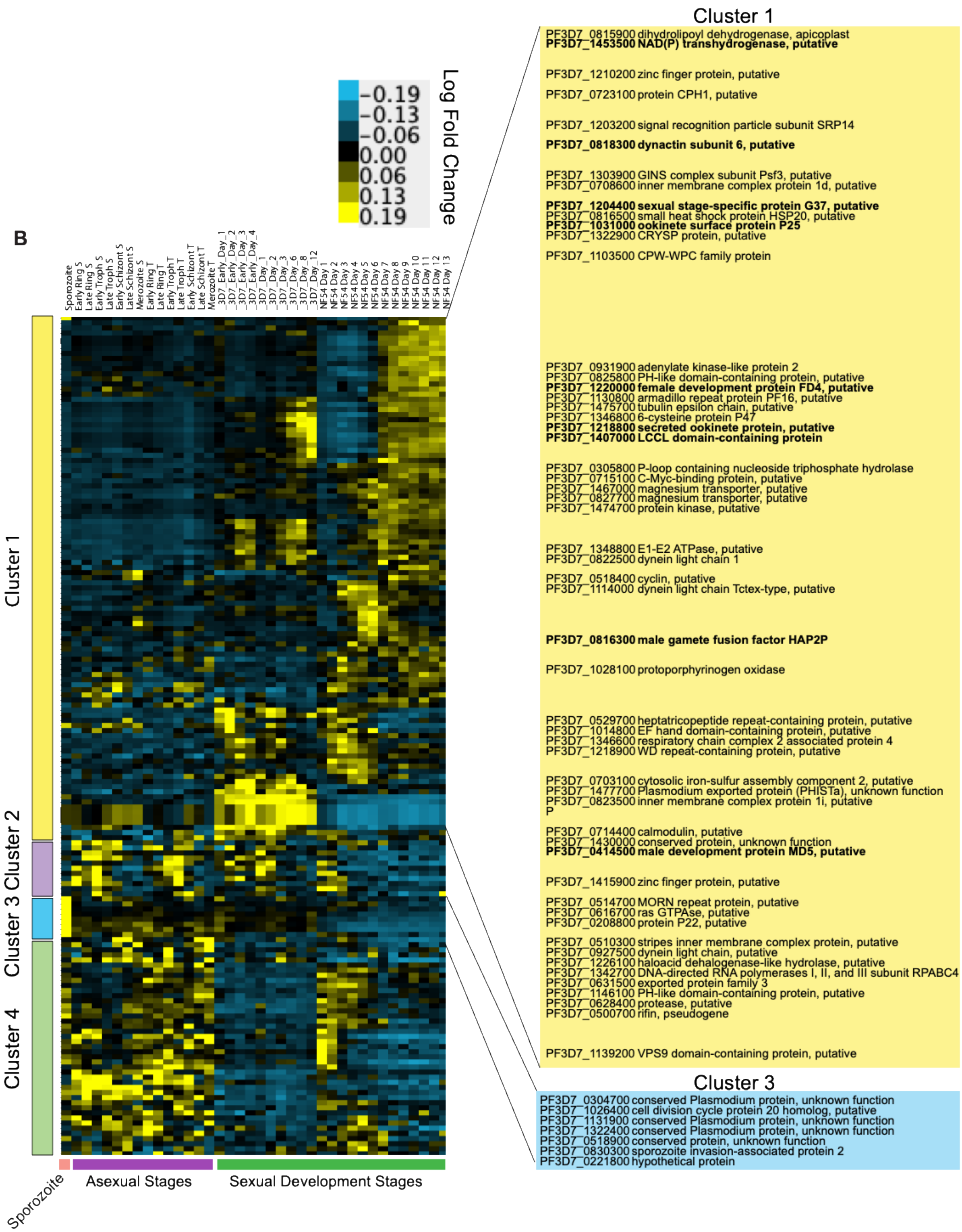

**Figure S3. Historical lifecycle expression for a subset mRNA transcripts upregulated in cryptosporin.** Lifecycle expression for 211 cryptosporin-upregulated transcripts was extracted from a published dataset (Young et al., 2005) for the 165 genes with a match and minimum expression of at least 50 units in one lifecycle condition. Historical lifecycle data was log-transformed, normalized and hierarchically clustered. For the historical data, purified asexual and sexual development stages had been obtained and analyzed using a quantitative microarray-based analysis (Young et al., 2005) with the stages shown on the top and bottom of the heatmap. Four clusters were identified using a correlation metric. Genes names with a predicted function for the clusters are shown in yellow (Cluster 1, mostly sexual development) and blue (Cluster 3 sporozoite). Genes shown in bold have known roles in parasite sexual development. The complete set of genes in the cryptosporin RNAseq study and in the clusters is given in Tables S1 and S2-4. The historical data, matched to modern gene names is given in Table S2.

### Supplementary Tables

**Table S1. Genes expressed above background after rRNA depletion and library-size normalization.** Attached Excel spreadsheet.

**Table S2. The 165 genes that showed ≥2-fold upregulation in cryptosporin were selected for plotting and clustering the lifecycle expression.** Attached Excel spreadsheet.

**Table S3. SNV/indel variants in cryptosporin-resistant clones (10 months after start of selection).** Attached Excel spreadsheet.

**Table S4. SNV/indel variants in cryptosporin-resistant flasks (5 months after the start of selection).** Attached Excel spreadsheet.

**Table S5. Copy number variants from whole genome sequencing analysis of cryptosporin-evolved bulk culture and clone samples.** Attached Excel spreadsheet.

**Table S6. Details of *Plasmodium* strains used in this study.**

| Parasite lines | Background | Method | Gene modified | Mutation | Reference |
|---|---|---|---|---|---|
| PfDd2 | Dd2 | clinical isolate | NA | NA | (Volkman et al., 2007) |
| Pf3D7 | 3D7 | clinical isolate | NA | NA | (Gardner et al., 2002) |
| *Pb*Luc | *Plasmodium berghei* | Double-crossover recombination | rRNA locus | Insertion of firefly luciferase | (Franke-Fayard et al., 2006) |
| PfPI4K_S1320L | Dd2 | zinc-finger nucleases | *pfpi4k* | S1320L | (McNamara et al., 2013) |
| PfCARL_I1139K | Dd2 | CRISPR/Cas9 | *pfcarl* | I1139K | (LaMonte et al., 2016) |
| PfACS_A597V | Dd2 | CRISPR/Cas9 | *pfacs* | A597V | (Summers et al., 2022) |
| attB | Dd2 | Single-crossover recombination | pfcg6 | attB site insertion | (Nkrumah et al., 2006) |
| attB::ScDHODH | Dd2 | Bxb1 integrase-mediated system | pfcg6 | ScDHODH insertion | (Painter et al., 2007) |

| | | | | | |
|---|---|---|---|---|---|
| 1-B7 | Dd2 | *in vitro* evolution | *pfaqp, pfsod-1, pfsod-2* | F138Y, copy number increase | this paper |
| 1-C8 | Dd2 | *in vitro* evolution | *pfaqp, pfsod-2* | F138Y, copy number increase | this paper |
| 2-F7 | Dd2 | *in vitro* evolution | *pfaqp, pfsod-1, pfsod-2* | F138Y, copy number increase | this paper |
| 2-F10 | Dd2 | *in vitro* evolution | *pfaqp, pfsod-1, pfsod-2* | F138Y, copy number increase | this paper |
| 3-D8 | Dd2 | *in vitro* evolution | *pfaqp, pfsod-2* | F138Y, copy number increase | this paper |
| 3-F5 | Dd2 | *in vitro* evolution | *pfaqp, pfsod-2* | F138Y, copy number increase | this paper |
| $2E2^{F138Y}$ | Dd2 | CRISPR/Cas9 | *pfaqp* | F138Y | this paper |
| $2E3^{F138Y}$ | Dd2 | CRISPR/Cas9 | *pfaqp* | F138Y | this paper |

**Table S7. Primers used in this study. Primers used for PT-qPCR in Figure 2S were listed as well as those used in CRISPR/Cas9 and overexpression studies.**

| Gene or Transcript ID | Type of research | Forward Primer(5'-3') | Reverse Primer(5'-3') |
|---|---|---|---|
| NTH | RT-qPCR | GATCGACGCGTGCACAAA | TGCCTGTAAACCTGCAACAC |
| NUP335 | RT-qPCR | ACAAAGATAACTTCCAAGGCA | ATCGAATTGGATTCTCCCAAA |
| PF3D7_0102900 | RT-qPCR | TGGCTACTTAGCACAATCCC | TGCTCGAAATACTGGACCAAC |
| ATPase1 | RT-qPCR | CCGGATGTCAGTGCATTATGT | TTCCACATTTCTTTGCGACA |
| TMK | RT-qPCR | ACGTGGTGTATGAACCCAGA | TCTTCATGTGCGAAGTGTTT |
| MFS6 | RT-qPCR | ACTCATACAGAGCACAGGAA | TGATGGTAGGAATTATTGGTTCT |

| | | | |
|---|---|---|---|
| MFS1 | RT-qPCR | GTGGCGCTAACGAATCACTT | TGTCATTTGGCTAGTTCCACA |
| PfAQP | PCR for CRISPR/Cas 9 | GAACTATATCACAATGCATA TGTTATT | GTGTATTACAAATCTACACC AT CTT |
| PkSOD-2 | Overexpressi on of PkSOD-2 | TATATATCCAATGGCCCCTT TCCGGGCGCGCCTAGAAGG TAATACGAAGTAGGATCATC | GCATAATCAGGTACGTCATA AGGGTATCCGGAGACGTCAT TTAATGTGGACAAGTTGTAG TTGG |
